# Predicting Trends in Avian Wing Motion and Energetics Across Scales

**DOI:** 10.1101/2025.06.10.658879

**Authors:** Ben Parslew

## Abstract

Birds exhibit monotonic trends in their flight kinematics across a broad range of scales. For example, in 10g to 10kg birds flight speed increases with scale while wingbeat frequency decreases. Existing hypotheses propose that aerodynamic and biomechanical factors may drive these trends. Predictive simulation models have used these drivers to computationally synthesize flying birds. However, only a small number of species have been simulated and so this approach has not explained the kinematic trends observed across different scales of birds.

This work uses predictive models to synthesize flight across all scales and shows that profile drag of the avian aerofoil is the key physical driver of self-selected kinematics. Profile drag causes the monotonic increase in flapping speed across scales and the near-constant ratio of flapping speed to flight speed (Strouhal number). The most efficient kinematics minimize Strouhal number to minimize the cost of transport, with the minimum cost of transport being bounded by the aerofoil maximum lift-to-drag ratio. The decrease of wingbeat frequency and increase of wing elevation amplitude with scale are attributable to balancing induced drag and the lateral component of profile drag.

The analytical and low-order numerical models presented provide transparency in understanding how modelling choices influence flight predictions. The work provides a platform for developing higher fidelity flight energetics models that address the scale limits of flying birds.

## Main

Avian flight energetics models can explain how mechanical and metabolic power consumption vary for different flight speeds and for different species^1^. These models input wing kinematic data and flight speeds from experimental measurements into mathematical flight physics models. This allows power- or energy-optimal flight speeds to be identified, for example^1,2^. Their core utility is to help explain the self-selected flight speeds of real birds. An alternative class of models known as *predictive models* aim to simulate flight physics from first principles, relaxing the requirement to input kinematics from experiments. These models were originally inspired by predictive models of human locomotion which have been instrumental in understanding gait and explaining why specific limb kinematics are preferred over others^3^.

Predictive models consist of a physics-based description of an animal that can automatically trial different locomotion techniques and identify those with preferable characteristics - such as minimal power or, joint torque. So the preferred motion is learnt, rather than prescribed. Pennycuick’s model^1^ is arguably the foundational predictive model of avian flight. It predicts the single variable of flight speed from physical properties of the bird and its environment. It has been used to predict how flight speed varies across different scales of birds by identifying flight speeds that require minimum flight power and minimum cost of transport (COT) (Fig. 1b). The advantage of such a minimalistic model is that its predictions are easily relatable to the model formulation and underlying physics – the equations predict that bird flight speed scales proportionally with mass^1/2^, and the main driver of this trend is that as bird scale increases wingspan does not increase as quickly as mass. However, this minimal model cannot predict the choice of wing motions, relying instead on empirical kinematic inputs.

**Fig. 1.**
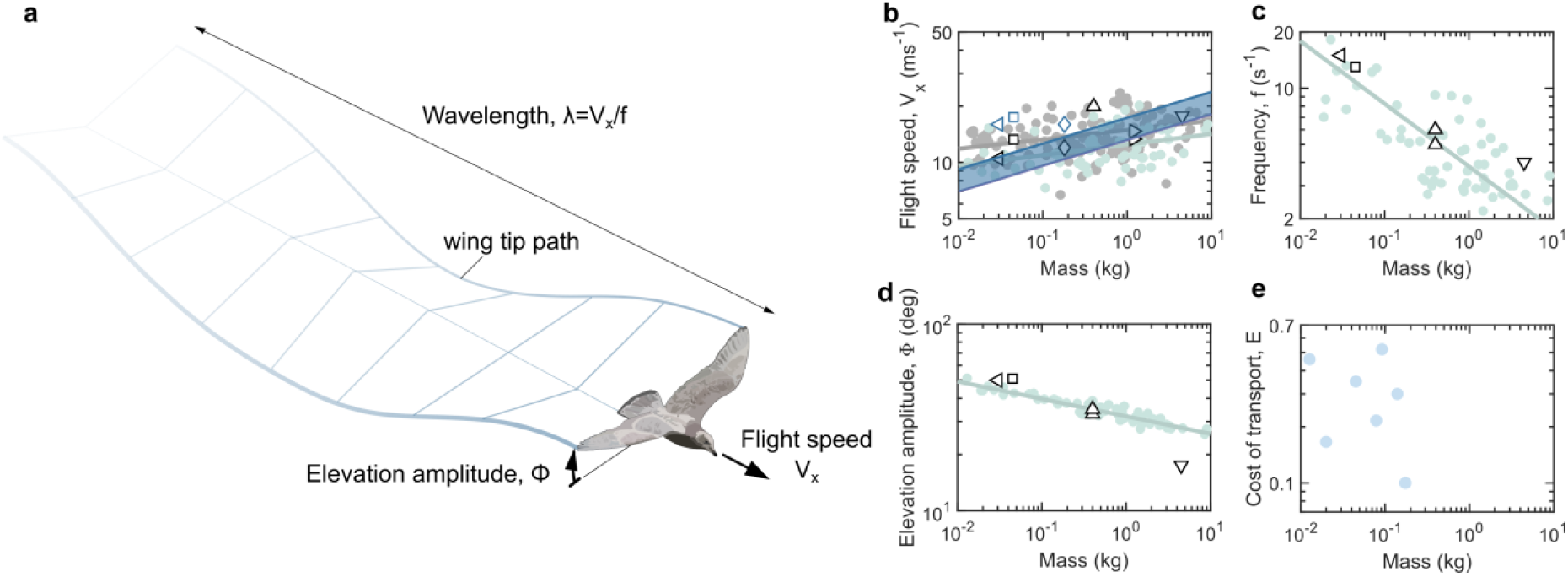
Kinematics and energetics across the scales of avian flight. **a** Diagram of fundamental cruising flight kinematic variables of flight speed, *V*_*x*_, elevation amplitude, Φ, wavelength, *λ* and frequency, *f*. **b-d**, Kinematic data for birds across scales from experiments (green symbols^10^, grey symbols^11^ with regression lines computed from nonlinear least squares regression using the Matlab 2023a function *“fit”*), from numerical predictive simulation models (black open symbols are from models detailed in Extended Fig. 1 for the following species: thrush^5^ - left pointing triangle; budgerigar^5^ – square; jackdaw^8^ – diamond; pigeon^6^ – upward point triangle; ibis^9^ – right pointing triangle; generic “shorebird”^4^ – downward point triangle) and from analytical predictive simulation model^1^ (blue open symbols correspond to species and references from the same black open symbols; blue shaded region bounded by the minimum power flight speed and minimum COT flight speeds predicted for allometrically scaled bird models^1^).**e** COT data collected from different species (see Method).

At the other end of the spectrum, predictive models of flight have synthesized both flight speed and wing motion using higher fidelity physics models and numerical optimization methods (Fig. 1)^4–9^. These use species-specific physical data. As different aerodynamic and inertial models are used for each species the findings do not allow the trends across different scales to be interpreted in relation to the underlying models (Fig1 b-d x). Also, their higher complexity reduces their transparency compared to earlier models^1^.

A new class of predictive model could find an effective compromise between these two approaches – retaining the transparency of semi-empirical predictive models while being capable of explaining the selection of wing kinematic across all scales of birds.

### Optimal flapping is optimal gliding

To understand why avian wing motions vary across scales this work firstly developed a minimal predictive flight model that incorporated wing kinematics from the outset. The baseline model was a vertically plunging wing with infinite aspect ratio, neglecting wing rotational effects and induced drag. The wing oscillates vertically with constant speed. The kinematics are characterized by the forward flight speed (*V*_*x*_) and the flapping speed (*V*_*y*_), or equivalently the plunging angle (*β*, Fig. 2 a,b). The avian body was initially neglected, negating parasitic drag. Therefore the power expenditure and cost of transport (COT) depend only on aerofoil profile drag. This was the simplest description of avian flight physics that retained information about wing kinematics; a similar approach was used to explore power-minimizing wing motions in hovering and forward flight^12^. The simplicity provided transparency that allowed easier interrogation the higher fidelity models introduced later.

**Fig. 2.**
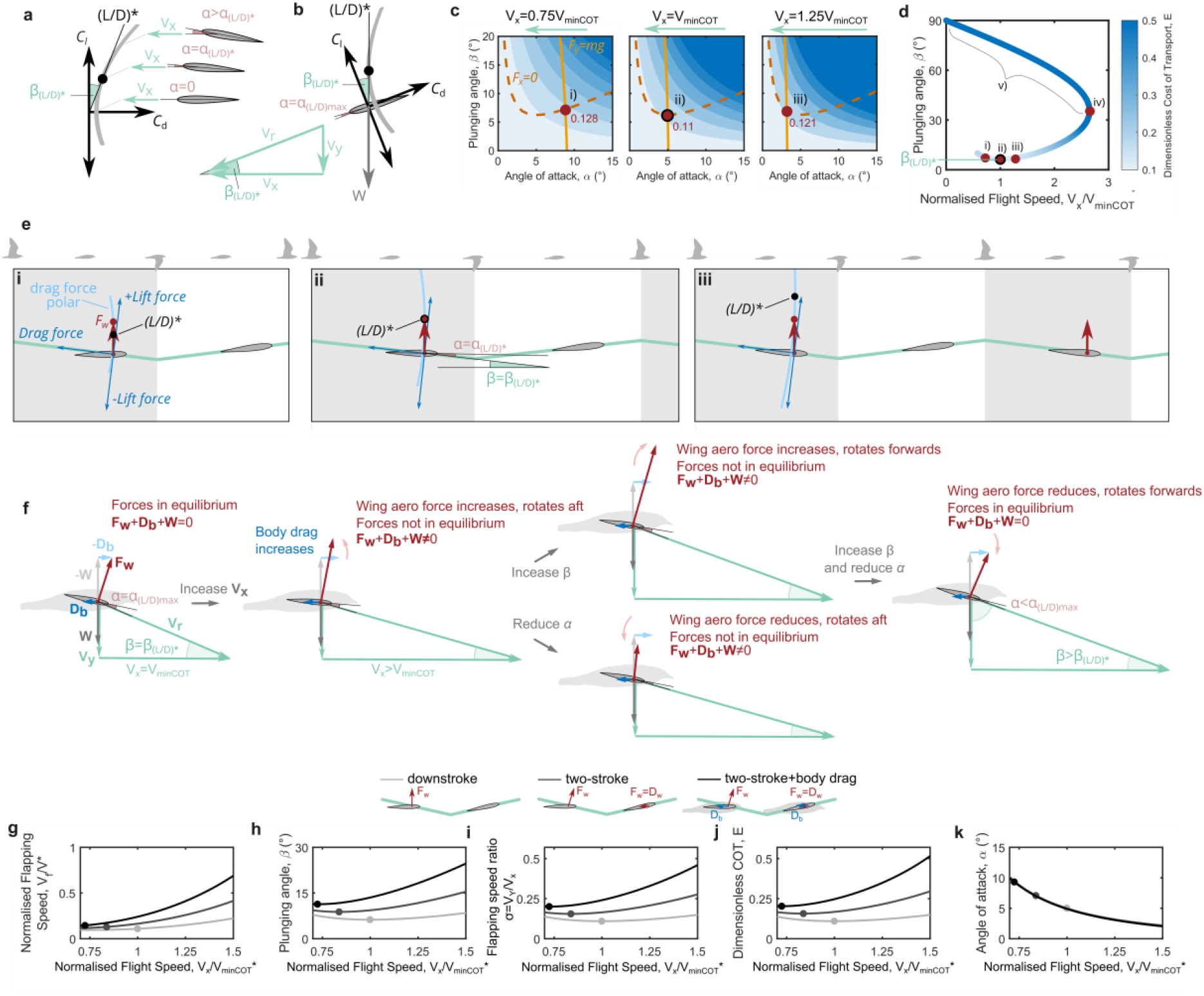
Seeking flight equilibrium through variations in wing kinematics and flight speed. **a** Illustration of a drag polar with the tangent representing the maximum lift/drag ratio (maximum glide ratio) and the angle between the tangent and the *C*_*L*_ axis being the plunge angle of a plunging wing for maximum lift/drag ratio. **b** The drag polar rotated to align with plunging wing in forward flight; at the angle of attack and plunge angle of maximum lift/drag ratio the aerodynamic force is parallel and opposite to the weight vector. **c** Predicted COT for the downstroke model for varying angle of attack, plunging angle and flight speed. The lines for vertical equilibrium (light orange) and horizontal equilibrium (dark orange dashed) intersect at the cruise condition (red dots). Point ii) is the minimum achievable COT of a plunging wing (from equation (5)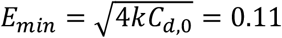; see Method). **d** Predicted COT-optimal kinematic solutions for cruising flight at different speeds, showing COT-optimal plunging angle as a function of flight speed. Angle of attack reduces from (i) through to (v). Typical flights would involve normalized flight speeds around 1. Solution (iv) is the maximum attainable flight speed of the downstroke model (extended equation 47). Solutions (v) use drag-based weight support. **e** Depictions of kinematics from solutions (i)-(iii) in **c** and **d**; polars of drag and lift forces (not coefficients; light blue lines) show that in COT-optimal flight (ii) the tip of the wing aerodynamic force vector (red dot) is coincident with the polar at the maximum L/D ratio. **f** Cartoon illustrating how from typical cruising flight conditions an increase in flight speed requires a combined increase in plunging angle and reduction in angle of attack to retain equilibrium. **g-h** Predicted kinematics for varying flight speeds for three different plunging models; flight speeds are normalized by the minimum COT flight speed for the downstroke model, *V*_*minCOT*_^*\**^. Dots indicate the COT-optimal solutions from analytical models (Supplementary Information) and lines are the solutions with varying speed from numerical models (Methods).

By neglecting the aerodynamics of the upstroke, the resulting *downstroke model* provided simple analytical predictions for wing kinematics that yield the lowest COT:

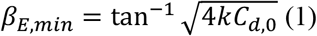

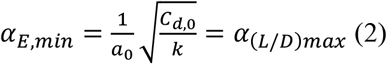

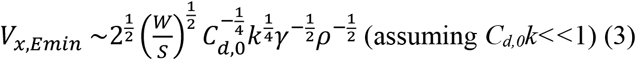

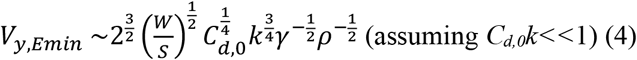

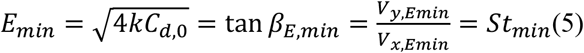

where *β*_*E,min*_, *α*_*E,min*_, *V*_*x,Emin*_ and *V*_*y,Emin*_ are the plunging angle, angle of attack, flapping speed and flight speed for the lowest COT, respectively, and *E*_*min*_ and *St*_*min*_ are the lowest COT and Strouhal number, respectively. The ratio of the durations of the downstroke to upstroke is the downstroke fraction, *γ*. The aerofoil drag is characterized by *k* and *C*_*d,0*_ – the lift-dependent and lift-independent viscous drag coefficients, respectively – while the aerofoil lift is characterized by the gradient of the lift curve slope – *a*_*0*_. The weight, *W*, and wing planform area, *S*, define the system scale. *ρ* is the local air density.

These equations reveal how the system physical variables influence the selection of kinematics. For example, COT-optimal flight speed and flapping speed are proportional to (wing loading)^1/2^ (equations 3 and 4) so if wing loading increases birds would be expected to flap and fly more quickly, which is what is observed in field studies^13–15^. The minimum COT is achieved by selecting the wing angle of attack for the highest lift/drag ratio (equation 5), and the prediction is independent of the downstroke fraction and scale of the bird. This result is identical to the condition for maximum range of a gliding fixed-wing aircraft^16^, which is unsurprising given that that the downstroke model treats the wing as a lifting surface undertaking intermittent glides with constant rates of descent.

The plunging angle required for minimum COT (equation 1) is also independent of downstroke fraction and scale and depends only on the lift-independent and lift-dependent viscous drag coefficients. An intuitive view of the plunging angle required for minimum COT is shown in Fig. 2a,b; the drag polar is rotated such that the tangent representing maximum lift/drag ratio is coincident with the weight vector. Taking this finding further, Fig. 2eii depicts to scale the COT-optimal kinematics that achieve equilibrium cruising flight and how this occurs at the maximum lift/drag condition; note that here the polar of drag and lift forces is plotted, rather than force coefficients. At lower and higher flight speeds force balance is only attainable using higher plunging angles, and higher and lower angles of attack, respectively, which increases cost of transport. (Fig 2ei, 2eiii, respectively)

Around cruise conditions the vertical force balance (*F*_*y*_=*mg*) is most sensitive to changes in angle of attack, whereas the horizontal force balance (*F*_*x*_=0) is most sensitive to changes in plunging angle (Fig 2c). This highlights the control variables that could be exploited by birds to elicit vertical and horizontal accelerations for maneuvering, or in response to gusts, for example. COT is similarly sensitive to angle of attack and plunging angle. The lowest COT of 0.11 (Fig2cii) represents a theoretical lower limit for all birds and is independent of scale. Induced drag, body parasite drag, and 3d wing flapping effects all lead to higher COT, which will be explored later.

By neglecting forces on the upstroke the downstroke model predicts some more nuanced solutions in other conditions. The model predicts that there is a maximum attainable forward flight speed (Fig 2 div). Attempts to increase flight speed further through adjusting the plunging angle or angle of attack upset the force equilibrium required for cruise. So to fly at higher speeds either the upstroke must be utilized or the angle of attack during the downstroke must not remain constant. Alternatively, high average flight speeds can be achieved using a series of downstrokes that periodically accelerate, decelerate, climb and descend, averaging to level flight over time. While this speed range is beyond that exhibited by birds, the finding is relevant to producing engineered flapping wing systems for high speed flight. At lower flight speeds a secondary set of wing motions exist with higher plunging angles and lower angles of attack, where drag is the primary weight support mechanism (fig 2cv). While birds do use drag-based weight support during short commutes^17^ typical cruising flight is characterized by lift-based weight support. However, it is noteworthy that such a simple model is able to capture these two distinct weight-support mechanisms.

### Efficient flight reduces Strouhal number

More representative models of avian flight were developed by including upstroke drag (*two-stroke* model) and body drag (*two-stroke-body* model). An initial graphical examination of the aerodynamic forces showed how a change in flight speed alone upsets both horizontal and vertical equilibrium in these models (Fig2 f). The plunging angle and angle of attack must both be adjusted to retain equilibrium with changing flight speed^18^. Flying faster than the COT-optimal speed must drive the angle of attack below its optimal lift/drag condition, and the plunging angle above it. So the assumption that energy-optimal flight involves the wing operating at its maximum lift/drag ratio can no longer hold when upstroke and body drag are considered^11^.

To quantify these predictions the downstroke model was extended to incorporate drag on the upstroke and on the body (Supplementary Information; Fig. 2g-k). The resulting solutions are closed form analytical expressions with only slight modifications to equations 1-5, and as such are intuitive for understanding how flight kinematics and COT vary with scale. To verify the analytical models a numerical solution was also obtained using computational optimization of COT (Methods; Fig. 2 g-k).

The additional lift-independent viscous drag from the upstroke and body increases COT at all flight speeds, and nearly doubles the minimum COT in comparison the downstroke model (Fig. 2j). This highlights the energetic incentive for birds to minimize upstroke drag (e.g. through wing flexion) and to minimize body drag (e.g. through having slender bodies). Extra lift-independent viscous drag reduces the COT-optimal flight speed. So if a bird were to increase body or wing drag coefficients, such as through injury or weight gain, there would be an energetic preference for slower flights.

The analytical models showed that in minimum COT flight the COT is equal to flapping speed ratio (equation 5). So to minimize COT, the flapping speed – or equivalently the plunging angle, Strouhal number, or reduced frequency – should be minimized, rather than targeted to a specifical optimal range. This finding aligns with wind tunnel measurements of engineered flapping wings. The practical significance of this finding is that Strouhal number or plunging angle are useful visual proxies for understanding changes in COT for an animal flying at different speeds, or for comparing differences in COT across different scales. This can be beneficial for comparing relative energetics during field studies, where kinematic data are more straightforward to record than power consumption.

### Drag dictates kinematic and COT limits

To understand how scale influences wing kinematics a conceptually simple starting point was to suppose that bird wing area scales proportionally to mass; this gives constant wing loading. The downstroke model then predicts scale independence of COT, and COT-optimal flight speed, flapping speed, flapping speed ratio and angle of attack (equations 1-5; extended Fig. 2a-e). The same scale independence was found to occur when wing upstroke drag and body drag are included, providing that body loading (weight/body frontal area) also remained constant (Fig. 3a-h). The addition of any sources of lift-independent drag (i.e. upstroke drag and body parasitic drag) simply reduced predicted flight speed and increased flapping speed but did not affect the scaling trends (Fig. 3 and extended Fig. 2). Extended Fig. 2f and 3a give graphical examples showing how doubling the animal weight does not affect the equilibrium of aerodynamic and gravitational forces for constant wing loading systems.

**Fig. 3.**
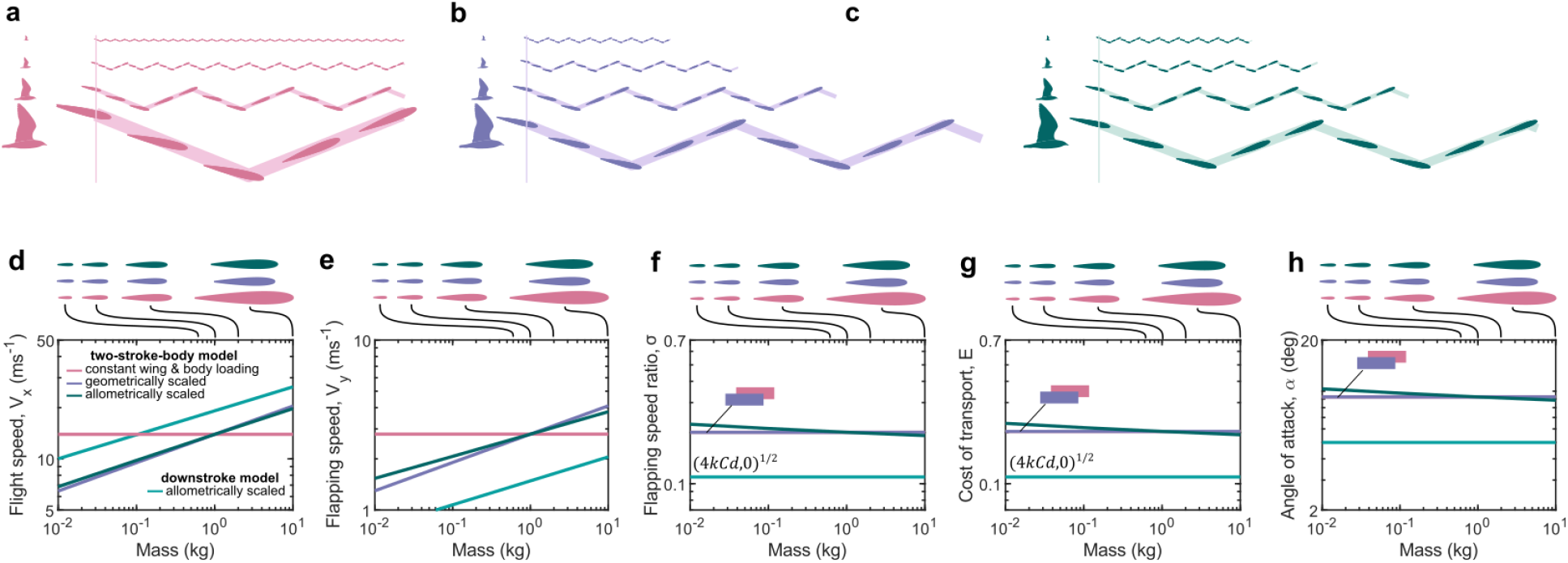
A comparison of different scaling models in predictive simulations of minimum COT cruising birds of 0.01-10kg. **a-c** Illustrations of changes in predicted wing kinematics across scales of birds (top to bottom, 0.01kg, 0.1kg, 1kg, 10kg) that have constant wing loading (**a**), are geometrically scaled (**b**) or are allometrically scaled (**c**). Illustrations assume the wing travels a vertical distance equal to the wing chord length so that changes in wing frequency can be depicted; this choice of distance is arbitrary as there are only unique solutions for flapping speed in these simulations, not for amplitude and frequency. All kinematics are shown over a fixed time window equal to the time period for a single wingbeat of the 10kg bird with constant wing loading (**a**). **d**-**e** Predicted COT-optimal kinematics for birds of different scales using different scaling models with the two-stroke-body model (pink, purple and green lines) and using the allometrically scaled downstroke model (turquoise) for reference as the ideal minimum COT case.

A second hypothetical system assumed wing and body area to scale geometrically with mass. This required different local wind speeds at different scales to balance weight and body drag. To achieve this the COT-optimal solutions varied flight speed proportionally with flapping speed across different scales (Fig. 3d,e). For example, doubling the weight of a geometrically scaled bird required flight speed and flapping speed to increase by a factor 2^1/6^ (Extended Fig 3b; equations 3-4). However, geometrically scaled systems retained the same constant flapping speed ratio, COT and angle of attack as constant wing loading systems (Fig 3f-h); the same is true of any system where wing and body loading scale proportionally. These kinematics retain the most energy-efficient plunge angle and angle of attack.

The allometric scaling of real birds sees wing area increasing faster than body area with increasing scale^1^. Incorporating this scaling into the models results in reductions in flapping speed ratio, COT and angle of attack with increasing scale (Fig. 3f-h; Supplementary Information Equations 58-68). However, the deviations of an allometrically scaled model from the simpler models mentioned previously are slight: across four orders of magnitude COT changes from 0.23 to 0.19 in allometrically scaled birds, and remains constant at 0.2 in constant wing loading and geometrically scaled birds. COT has a stronger dependance on bird wing and body drag coefficients than it does on their deviations from geometric scaling (equation 68). This finding can guide future experimental analysis to obtain more accurate models of wing and body coefficients, rather than more accurate records of wing and body shape, in order to make the biggest gains in understanding scaling trends in energetics in birds. Furthermore, improved understanding of lift independent and lift dependent drag coefficients of the avian aerofoil will help to better define the absolute lower limit of COT and Strouhal number for birds (equation 5; Fig. 3g, turquoise line).

### Lift-dependant drag controls wing speed

In high fidelity predictive models the influence of modelling decisions on the model predictions is not always evident. Simple models offer transparency in this area. The analytical models developed here showed the relationship between model input parameters and the predicted kinematics and COT. For example, wing flapping speed was predicted to vary monotonically with lift-independent and lift-dependant drag coefficients, however, flapping speed is more sensitive to changes in the latter (equation 4). Understanding these sensitivities guided us as to which input parameters should be defined more accurately when constructing a predictive model.

A sensitivity analysis was conducted of the two-stroke-body model to elicit the influence of the core aerodynamic inputs (Fig. 4, extended Fig. 4). Input variable ranges were defined using those from previous predictive models^4–9^ and from experimental measurements of avian wing and body aerodynamics ^14,20,21^. Predicted flight speeds were found to be similarly sensitive to the tested ranges of wing and body drag coefficients (Fig 4e, extended fig 4e,n). But flapping speeds and COT varied more across the range of tested lift-dependant viscous drag coefficient than across the range of lift-independent wing and body drag coefficients (fig 4g,h; extended Fig. 4g,h,p,q).

**Fig. 4.**
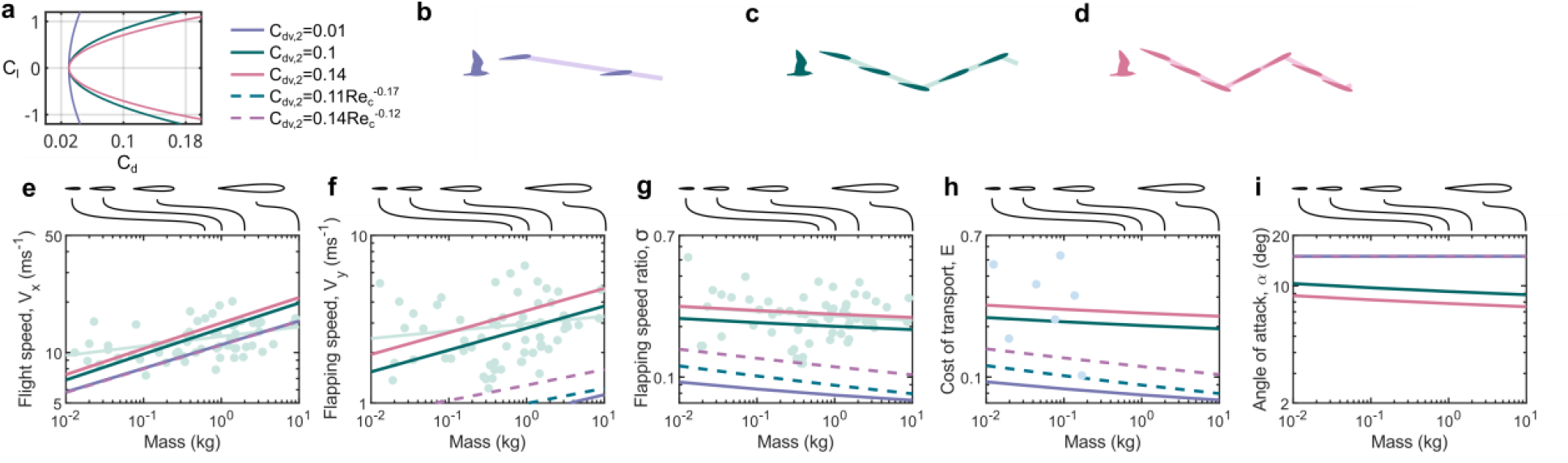
Sensitivity of the COT-optimal predicted kinematics to the lift-dependent viscous drag coefficient, *C*_*dv,2*_, using the two-stroke-body model. For constant drag coefficients the values tested were chosen based on the minimum value used in previous predictive simulations of bird flight^5^ (*C*_*dv,2*_=0.01), and data from wind tunnel tests on aerofoils at representative Reynolds numbers of flying birds (owl^21^: *C*_*dv,2*_=0.1; Eppler 387^20^ - *C*_*dv,2*_=0.14). **a** drag polars shown for three (constant) values of lift-dependent viscous drag coefficient. **b**-**d** illustrations of predicted kinematics for a 1kg bird; all kinematics shown over a fixed time window equal to the time period for a single wingbeat of the 1kg bird in Extended Fig. 4b. e-i predicted COT-optimal kinematics for birds of different scales using constant (solid lines) and Reynolds-number-dependent (dashed lines) models of lift-dependent viscous drag coefficient from previous predictive simulations^5^

Interestingly, the predictions of kinematics and COT were insensitive to the specific flapping kinematics, with constant and sinusoidal flapping speeds yielding similar results (extended Fig. 4s-y). So future predictive models would benefit more from accurate models of avian aerofoil lift dependant viscous drag coefficients than they would by attempting to capture more intricate wingbeat kinematics.

### Birds trade-off lateral and induced drag

A plunging wing does not distinguish between wingbeat amplitude and frequency. This is illustrated in Extended Fig 5a,b where equal flapping speeds but different amplitude and frequency combinations are shown to generate the same aerodynamic force and power; Extended Fig. 5c shows that an infinite number of amplitude and frequency combinations can achieve equilibrium cruise conditions with the same COT. This limits the scope for predicting kinematics of avian flight using plunging models. To rectify this, rotating 3d wings were modelled numerically to capture the elevation and depression of the avian shoulder joint. However, moving from 2d plunging to 3d rotating involved two distinct changes to the system physics which both increased COT, and which each drove the wing kinematics to opposing preferred solutions.

Firstly, rotating the wing resolves a component of the aerodynamic lift laterally. This requires additional flapping speed for weight support, which increases the lateral profile drag and COT (extended Fig. 6a-f). The preferred solution tends to zero rotation amplitude and infinite frequency – equivalent to 2d plunging. Secondly, the 3d wing generates a tip vortex which creates downwash and induced drag, and increases COT. Sweeping the wings over a greater area reduces the induced drag, so the preferred solution tends to an infinite rotation amplitude and zero frequency. Birds have to find a compromise between these two energy sinks.

3d wing flapping effects were explored by adding wing rotation and a simple actuator disk model of induced velocity to the two-stroke-body model (Fig. 5). Including induced velocity enables minimum power and minimum COT flight to be predicted; otherwise the minimum power solution is trivial, tending to the maximum angle of attack constraint and lowest achievable flight speed. The minimum power and minimum COT speeds are likely to envelope the self-selected flight regimes occupied by birds in nature^1^ making the model representative of experimental field study conditions.

**Fig. 5.**
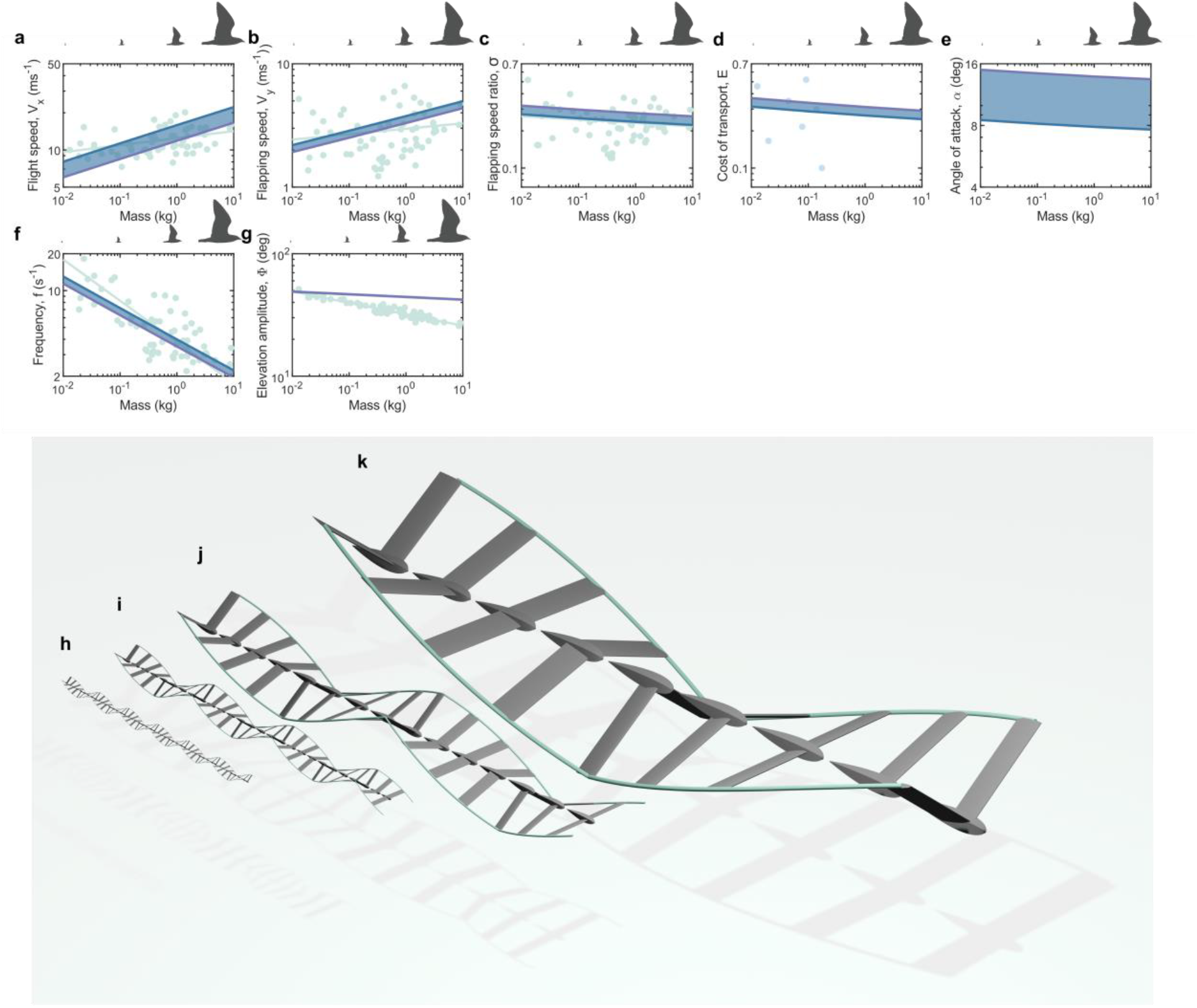
Predicted trends of flight speed, COT and 3d wing motion across scales of birds. **a-g** Predicted kinematics for COT-optimal (blue line) and power-optimal (purple line) flight, with self-selected kinematics expected to lie between these two (shaded blue region). **h-k** Visualizations of predicted kinematics for COT-optimal flight across for birds of mass 0.01, 0.1, 1 and 10kg, illustrate how bird kinematics might be viewed during field observations. All cases plotted of equal time windows equal to the time period of a single wingbeat of the 10 kg bird

The model predicted a slight monotonic reduction in elevation amplitude with increasing scale. Greater reductions in elevation amplitude were seen in field studies^10,22^ (Fig. 5g). However, it should be noted that experimental definitions of this angle differ among the data sets used and are complicated by deformation of the wing (see methods). It is noteworthy that the elevation amplitudes for minimum power and minimum COT only differ by less than 1%. For this reason, elevation amplitude is not a useful visual proxy during field study observations to infer whether birds are flying at minimum power, minimum COT, or some other characteristic speed. This is illustrated through simple 3d visualisations (Fig. 5h-k) where it is challenging to identify even the larger changes in elevation amplitude that occur between different scales of birds, let alone the differences in amplitude between minimum power and minimum COT flight. Clearer to observe is the increase in flight speed with increasing scale, with larger birds covering a greater distance than smaller birds over a fixed time window. Predicted trends in flight speed were similar to those from earlier analytical models^1^ (Fig. 1b, Fig. 5a), with both predicting a sharper trend than that recorded in field observations. A limited number of species have been predicted from numerical species-specific models^4–6,8^. However, these predictions suggest that species-specific input parameters may lead to higher flight speed predictions for smaller scale birds than those from the present model, and better capture the trends in flight speeds seen in experiments.

Notably, the model predicted the sizeable reduction in wingbeat frequency seen in experiments of almost an order of magnitude over the four orders of magnitude of bird scale (Fig. 5f). The selected wingbeat frequency was found to be governed by the balance of lateral profile drag and induced drag. So a quasi-steady aerodynamic model of a 3d flapping wing and body can predict wingbeat frequencies of birds using only allometric expressions for wing and body area scaling, and without requiring kinematic data from other birds. This extends previous methods of frequency scaling which rely on empirical kinematic data^15,23^.

A narrow range of flapping speed ratios was predicted between power-optimal and COT-optimal flight across all scales and this captures well the trends seen in field experimental (Fig. 5c). This provides evidence that the tendency of birds to utilise specific Strouhal numbers is not an effort to harness unsteady aerodynamics benefits^24^ – which are not modelled here – rather, birds simply select the lowest possible Strouhal number that can sustain the equilibrium of forces for cruising flight^18,19^. This finding also has pragmatic applications for flight research: using the trace of the wingtip path as a guide (Fig. 5h-k), observing the steepness of the wingtip path of a flying bird is a visual cue of their closeness to optimal flight conditions.

Minimum-power flight was predicted to use angles of attack close to typical stall angles (Fig. 5e). While birds can likely manage stalled flow conditions more effectively than engineered aircraft can, it is still undesirable^25^. So birds are likely to avoid flying at minimum power speed unless absolutely necessary^1^. As flight speed increases from minimum power to minimum COT speed the reduction in angle of attack is sizeable – from around 15° to 8°. Even through nominal angle of attack is challenging to capture experimentally, it is viable to observe changes in angle of attack of this magnitude. This means that transitions between characteristic minimum power and minimum COT states may be identified from angle of attack measurements alone.

The model predictions were found to be robust to key aerodynamic modelling decisions. Predicted kinematics and COT were relatively insensitive to the broad range of induced power factors that has been debated in literature^26^ (extended Fig. 7a-g), so assuming uniform downwash (*k*=1) is a simple and insightful initial approximation. There was limited difference in predicted kinematics between models that assumed a uniform chord distribution and ones that used an avian wing chord distribution (extended Fig. 7h-n). Fundamental insights into the selection of kinematics across all scales of birds can be obtained using low order aerodynamic and wing geometric models.

### Flight cost fractions change with scale

The cost of transport was broken down into its constituents to observe their influence across different scale (Fig. 6a,b). These energy sinks reduce the overall efficiency – defined as effective lift/drag ratio of the bird^1^ *-* in comparison to the baseline ideal system of a 2d aerofoil (Fig. 6c). The relative impacts of the sinks vary across scales. For example, parasitic drag accounts for 22% of aerodynamic contribution to COT for the smallest birds but less than 15% for the largest birds (Fig. 6a). So smaller birds have the most to gain from reducing their body drag coefficients. Conversely, upstroke drag contributes 15% aerodynamic COT to the smallest birds and 19% to the largest birds, so larger birds benefit more from reducing wing exposed area during the upstroke or through using the upstroke to generate lift.

**Fig. 6.**
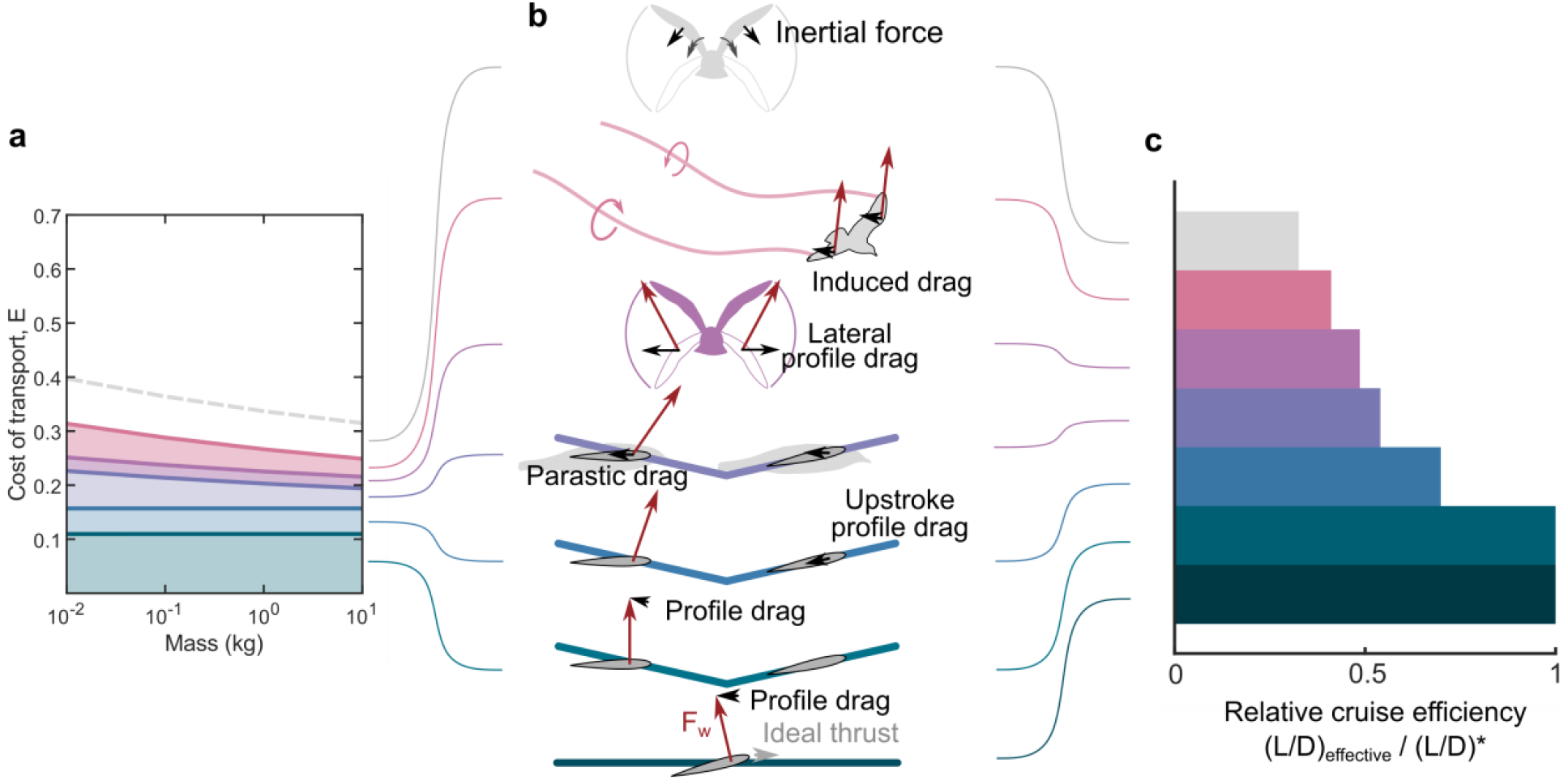
Energy sinks in cruising flight. a The cumulative sources of COT, with coloured regions indicating the fractions of COT attributable to aerodynamic energy sinks. The maximum additional energy that could be required to overcome inertial loads is indicated by the grey dashed line. **b** Cartoons of energy sinks, showing the aerodynamic force vector generated by the wing (red arrows) and the cumulative force components requiring energy consumption to overcome (black arrows). The baseline ideal case is shown as a fixed-wing of infinite aspect ratio with an ideal thruster to overcome profile drag only (green line). **c** The influence of the different energy sinks on the whole bird efficiency, defined as the ratio of the effective lift/drag ratio of the whole bird (equal to the reciprocal of COT) to the maximum lift/drag ratio of the aerofoil.

The inertial cost of accelerating and decelerating the wing is likely to be alleviated through elastic storage mechanisms^27^. However, a consistent mechanism has not yet been identified to describe the degree of energy storage across different scales. To illustrate the potential influence of inertia Fig. 6a explores the worst case scenario with no elastic recovery. Here, across all scales inertial effects would increase COT by over 25% compared to aerodynamic effects alone. So it can be hypothesized that across all scales birds are similarly likely to exhibit effective elastic energy storage mechanisms.

### Conclusions

In summary, the results revealed that low-order analytical and computational predictive models can be used to explain trends in wing kinematics across all scales of birds. Birds minimise their Strouhal numbers to minimise cost of transport. In treating the flapping wing as a simple 2d glider traditional aerodynamic theory explained the near-constancy of Strouhal number across the scales of birds. While extending to a 3d rotating wing model found that the wing elevation amplitude and frequencies are selected as a compromise between profile and induced drag on the wings, and that no empirical kinematic data are required to predict these trends.

The cost of transport was shown to be attributable to different energy sinks which have varying relative contributions across scales. Because of this, birds at different scales are likely to have evolved different morphological strategies for reducing cost. Solutions from this work can seed higher fidelity models that elucidate these morphological trends. In doing so, future models can exploit the fact that selected kinematics are most sensitive to the lift-dependant drag coefficient than to differences between scaling laws. This also helps to guide future experiments to collect complementary data that best improves model accuracy, and hence has the biggest impact to our understanding of kinematic trends across scales.

## Methods

### External Data Sources for Kinematics and COT

Kinematic data on flight speed, wingbeat frequency and wing elevation angle used for comparison to simulated values from the model (Fig. 4, 5, extended fig 4, extended Fig. 6) were taken from previous literature values, which are an accumulation of data from different sources^10,22^.

The flapping speed ratio, *σ*, was defined through the work as the ratio of normal flapping speed to forward flight speed (see Flight Mechanics Model). Applying the same definition to the experimental data gives

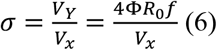

which assumes a constant flapping speed, a downstroke fraction of one half, and an aerodynamic control point located at *R*_0_ - the radius of the second moment of area (see Supplementary Methods). This makes the experimental and simulated data directly comparable. The flapping speed ratio and Strouhal number are related by

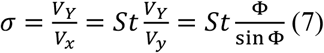

so as the wing rotation amplitude tends to zero (equivalent to a 2d plunging wing) the flapping speed ratio tends to the Strouhal number. Flapping speed ratio is used in the present study when making comparisons to experimental data, which capture wing rotation.

The experimental kinematics data is drawn from sources with different definitions of the elevation angle. For example Pennycuick^28^ defines the angle using photographs with chord lines drawn approximately tangential to the wing upper surface at the shoulder joint when viewed downstream of the bird; Scholey^29^ draws chord lines drawn on the proximal wing; Spedding^30^ takes measurements from measurements of the wake. Moreover, the use of a chord line involves assuming the wing as a single planar surface, so neglects details of wing bending and wrist circumduction. The experiments consist of both field experiments^29^ and wind tunnel experiments (e.g.Pennycuick^28^), where the latter was used to define average kinematic values over the range of tested wind tunnel speed^10,22^. Despite these factors, this remains the most comprehensive experimental data set on avian flight. A supplementary kinematic data set on flight speed was included in Fig. 1b taken from a standalone study using tracking radar^11^. Best fit lines of experimental data were obtained using nonlinear least squares fitting via the function *fit* in Matlab r2024b, using the trust-Region-reflective algorithm.

Calculations of flight speed from the previous predictive model^1^ (Fig. 1) used the equations for minimum power speed:

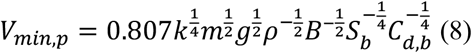

and maximum range speed

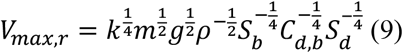

where the body reference area, *S*_*b*_, and actuator disk area, *S*_*d*_, were calculated using the approach from the same model, wingspan *B* was taken from least squares regression of the morphological data provide from the same reference, and the same reference values were used for induced power factor, *k*=1.2, and body drag coefficient, *C*_*d,b*_ = 0.1.

Experimental measurements of COT (Fig. 1) were taken from different sources using a range of measurement techniques to estimate mechanical power output: magpie, using bone strain measurements^31^; cockatiels and doves using bone strain measurements^32,33^; budgerigars and zebra finches using muscle activity and strain trajectory^34^, cockatiels using in vitro muscle measurements^35^; blackcaps using particle image velocimetry of the wake^36^. Techniques capturing metabolic COT were not included in the analysis.

### Flight Mechanics Model

The general flight mechanics equation used to describe cruising flight with a flapping wing were

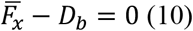

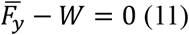

where 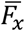 and 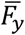 are the mean horizontal and vertical components of aerodynamic force on the wing, respectively, *D*_*b*_ is the body drag, and W is the weight (See supplementary Methods). These equations assume steady level flight of the bird, with constant flight speed throughout the wingbeat.

The mechanical power expenditure, dimensionless cost of transport and flapping speed ratio were calculated as

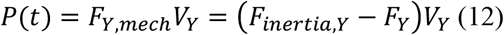

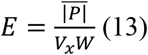

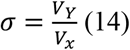

where *F*_*Y,mech*_ and *V*_*Y*_ are the mechanical force and wing flapping velocity components normal to the wing plane at the aerodynamic control point. Note that flapping speed ratio is the ratio of normal flapping speed to forward flight speed, whereas Strouhal number is the ratio of vertical flapping speed to forward flight speed (see supplementary methods); flapping speed ratio is the preferred term here as it is more general in capturing the ratio of characteristic speeds that influence the wing aerodynamics both for 2d plunging and 3d rotating wings. The mechanical force was obtained from the inertial force due to wing acceleration, *F*_*interia,Y*_, and the quasi-steady aerodynamic force, *F*_*Y*_; this is equivalent to calculating the power as the product of mechanical torque and angular velocity. For 2d plunging wing models *F*_*Y,mech*_ and *V*_*Y*_ are the vertical components of force and wing flapping velocity; for 3d rotating wings these components are resolved by the wing rotation angle, *ϕ* (Supplementary methods). 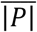 is the mean absolute mechanical power. This model assumes that energy is consumed for both positive and negative mechanical power. The definition of COT given here is the same as that used another recent work on avian aerodynamic efficiency^19^.

The aerodynamic force, *F*_*Y*_, was derived from the lift and drag on the wing. The method of resolving aerodynamic forces and velocities from local wind reference frames to other reference frames is detailed elsewhere^6^ so is omitted here for brevity. Lift and drag on the wing were modelled using a quasi-steady blade-element method, which neglected unsteady aerodynamics effects. The blade-element method models the wing as a series of 2d aerofoils of infinitesimal width along the wing from shoulder to tip, and neglects the influence of spanwise flow. The local aerodynamic force per unit wing length on the 2d aerofoils was integrated along the wing length to obtain the net instantaneous aerodynamic force on the entire wing (see supplementary methods).

The 2d aerofoil aerodynamic force per unit length is proportional to the local wind dynamic pressure, chord length, and local aerodynamic force coefficients, which were modelled here as

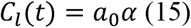

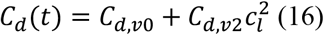

where *C*_*l*_ and *C*_*d*_ are the 2d aerofoil lift and drag coefficients respectively, *a*_0_ is the lift-curve slope, *α* is the aerofoil absolute angle of attack, *C*_*d,v*0_ is the lift independent viscous drag coefficient, and *C*_*d,v*2_ is the lift dependant viscous drag coefficient. The value of *a*_0_ = 2*π*rad^-1^ from thin aerofoil theory was used; this influenced the predicted angle of attack, but not the predicted wing kinematics. The use of a quadratic drag polar in equation (14) is known to neglect some of the aerodynamic details seen in aerofoils at avian-relevant Reynolds numbers ^18^, however this approach has proved fruitful in higher fidelity predictive models of avian flight^8^. The default value of *C*_*d,v*0_ = 0.03 was taken as an intermediate value from previous studies of avian aerofoils that estimated values from 0.02-0.045^21,37^. The value of *C*_*d,v*2_ = 0.1 was based on previous estimates 0.05-0.1^21^, though it should be noted that this value has been shown to be sensitive to aerofoil geometry^18^. As the model here aimed to be representative across all scales of birds rather than for specific species these generic values of *C*_*d,v*0_ and *C*_*d,v*2_ were used within a sensitivity analysis to explore their influence on kinematic predictions.

To calculate the time-dependant local aerodynamic force coefficients the angle of attack was calculated between the aerofoil zero lift line and the local wind velocity vector as

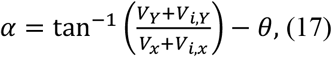

where *V*_*i,x*_ and *V*_*i,Y*_ are the components of induced velocity parallel to the flight direction and flapping velocity vector, respectively. *θ* is the aerofoil pronation angle-rotation about an axis along the wing length. The angle of attack, flight speed and flapping speed were independent variables in the model and the pronation angle was derived from equation (17). For the 2d plunging wing models and for 3d rotating wing models that neglected induced velocity, *V*_*i,x*_ and *V*_*i,Y*_ were zero. In other cases the total induced velocity was calculated using an actuator disk model with Glauert’s high speed approximation^38^ as

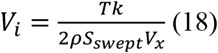

where *T* is the mean thrust over a single wingbeat, *k* is the induced power factor, and *S*_*swept*_ is the area over which the flapping wing is assumed to impart momentum to the fluid, which was modelled here as a circular sector defined by the wing flapping amplitude Φ and wing length, *l*_*w*_ ^6^:

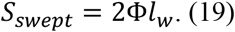

The orientation of the actuator sectors was defined relative to the horizontal plane by the angle

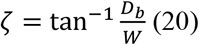

where *W* is the weight and *D*_*b*_ is the body drag. The body drag was given by

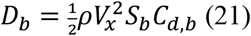

where *S*_*b*_ is the body reference area and *C*_*d,b*_ is the body drag coefficient. The thrust, *T*, which acts perpendicular to the sectors, was then given as

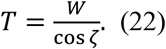

This allowed the induced velocity components in (17) to be calculated as

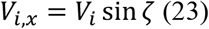

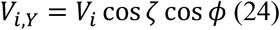

where *ϕ* is the rotation angle (elevation angle) of the wing about the shoulder joint measured from the horizontal plane; for plunging models *ϕ* = 0.

For a 3d wing rotating the angular acceleration, 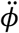, was used to calculate the inertial force in equation (12) as

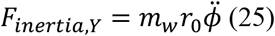

where *m*_*w*_ is the wing mass and is the wing radius of gyration.

A summary of the scaling expressions used to define geometric and inertial expressions is given in the Supplementary Methods.

### Kinematic Parameterization

Flight kinematics are defined using the flight speed, *V*_*x*_, and wing kinematics including the angle of attack, *α*(*t*) and flapping speed, *V*_*Y*_(*t*). For 2d plunging wing models the flapping speed is simply equal to the vertical velocity of the wing in the global axes

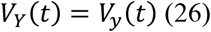

and for 3d rotating wing models the flapping speed is given as

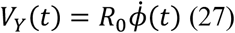

Where *R*_0_ is the radius of the second moment of area of the wing (see supplementary Methods).

Wing flapping kinematics were generally defined using the following piecewise function for the flapping velocity of the plunging models

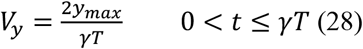

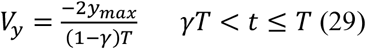

where *y*_*max*_ is the vertical displacement amplitude of the wing.

For the 3d rotating wing models wing flapping kinematics were generally defined using the following piecewise functions:

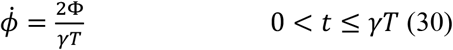

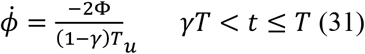

For the special cases where inertial effects were modelled (Fig. 6; extended Fig 4u-y) wing kinematics were defined using sinusoidal functions:

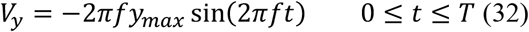

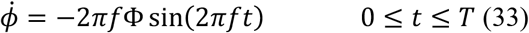

The angle of attack is defined using the following piecewise model for all cases:

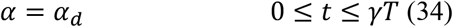

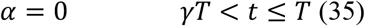

*α*_*d*_ is the downstroke angle of attack, shortened to *α* in the main text. Note that for the analytical downstroke model the force on the upstroke is neglected, whereas for all other models the upstroke force is modelled as the zero-lift drag.

### Analytical and Numerical Solutions

Analytical solutions of COT were obtained for the downstroke, two-stroke, and two-stroke-body models, which assume a vertically plunging wing with constant flapping speed and no induced velocity. Derivations of these solutions are given in the supplementary materials.

Numerical solutions were obtained using computational optimisation methods. This involved calculating the instantaneous force and velocity on the wing at *N* discrete times throughout the flapping cycle, and then calculating the mean absolute power as

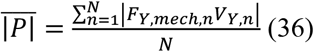

The output from equation (36) was then used in equation (13) to obtain the COT.

The minimum COT was obtained by numerically optimising the flight speed and wing kinematics. Formally, the optimisation problems are given here for plunging wing models as

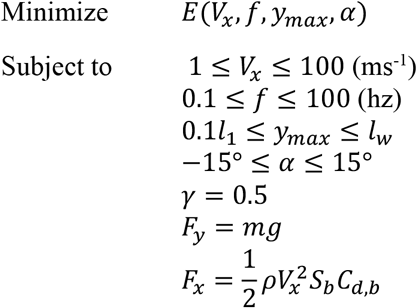

and for 3d rotating wing models as

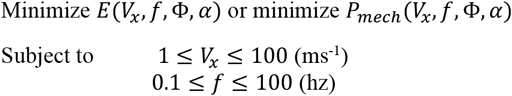

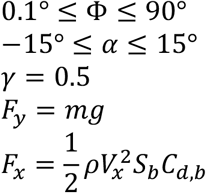

The equalities for *F*_*y*_ and *F*_*x*_ ensure that the force generated by the flapping wing balances weight, and body drag, respectively.

This formulation reduces the number of kinematic parameters as much as possible for transparency while still retaining enough information to furnish a predictive model of avian flapping flight. The downstroke fraction was constrained to *γ* = 0.5 for all numerical simulations presented here. Trials with different downstroke fractions found that an optimal downstroke fraction of around 0.55 across all scales of the rotating wing model reduced COT by less than 1% (Fig. 5), which is similar to that found in recent experiments on flapping wing models^19^. However, with this higher downstroke fraction the predicted kinematics changed consistently across scales, so the trends and conclusions from this work remained the same; predicted flight speed reduced less than 1%, frequency increased around 2%, elevation amplitude increased around 6%, and angle of attack increase by around 1%. As this did not influence the predicted trends the additional complexity was not warranted, so the simpler route was taken here of assuming that the upstroke and downstroke had equal durations.

The numerical optimisation problem is solved using the interior point algorithm in the Matlab 2023a “fmincon” function, which finds the minimum of a constrained nonlinear multivariable function. The constraint and optimality tolerances were both 10^−6^, and the step tolerance was 10^−10^. Each optimisation was initiated with multiple combinations of starting variables at their minimum and maximum constrained values, giving 16 combinations in total. The global solution was obtained as the solution with lowest COT from all starting locations; the majority of starting solutions that led to feasible solutions, led to the same feasible solution, with the minority of feasible solutions at much higher COTs.

## Extended figures

**Extended Fig. 1.**
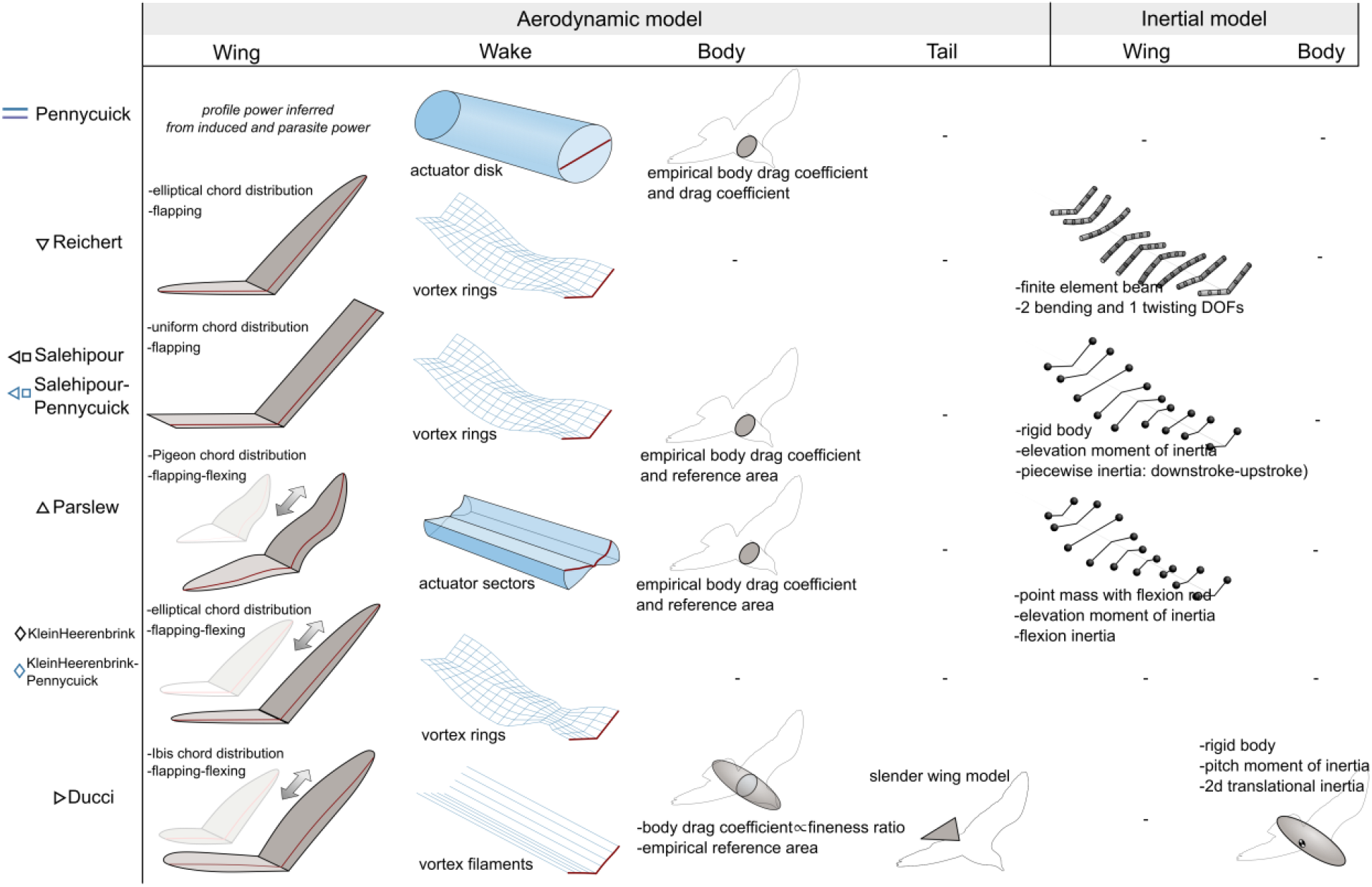
Pictorial representation of previous prediction simulation models of birds^1,4– 6,8,9^. Symbols and lines mirror those use in Fig. 1. “Selihipour-Pennycuick” and “KleinHeerenbrink-Pennycuick” are the use of the analytical predictive model of flight speed^1^ by authors within their own studies which develop numerical predictive models^5,25^

**Extended Fig. 2.**
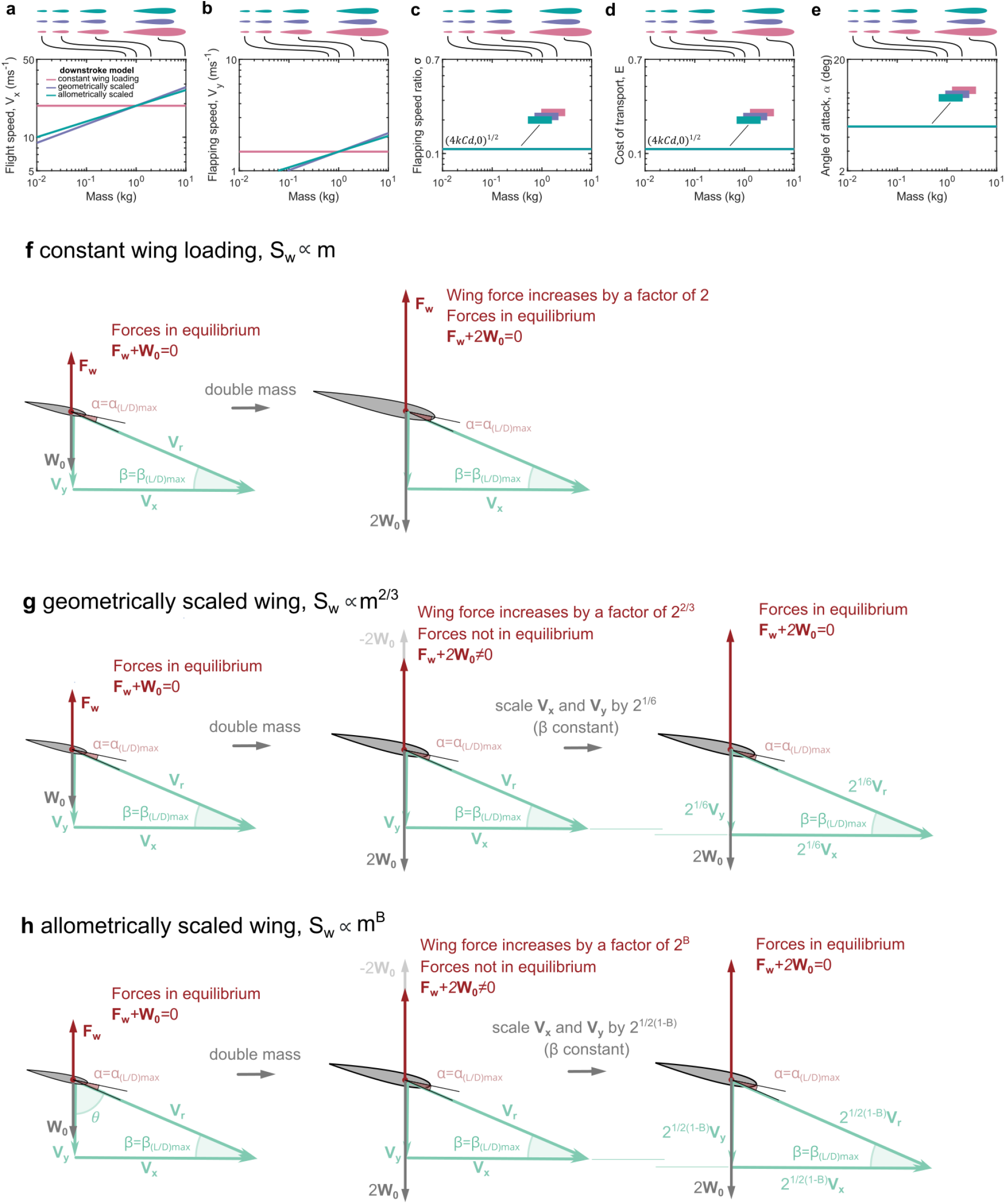
A comparison of different scaling models of wings in predictive simulations of minimum COT cruising flight. **a-e** Predicted COT-optimal kinematics for wings of different scales using different scaling models with the two-stroke model. **f-h** Cartoons illustrating an example of how doubling mass influences required kinematics for cruise depending on the scaling method: constant wing loading systems (**f**) require no changes to kinematics; geometrically scaled wings (**g**) require flight speed and flapping speed to scale by 2^1/6^; and allometrically scaled wings (**h**) require flight speed and flapping speed to scale by 2^1/2(1-B)^, where the wing area is defined by *S*_*w*_ ∝ *m*^*B*^.

**Extended Fig. 3.**
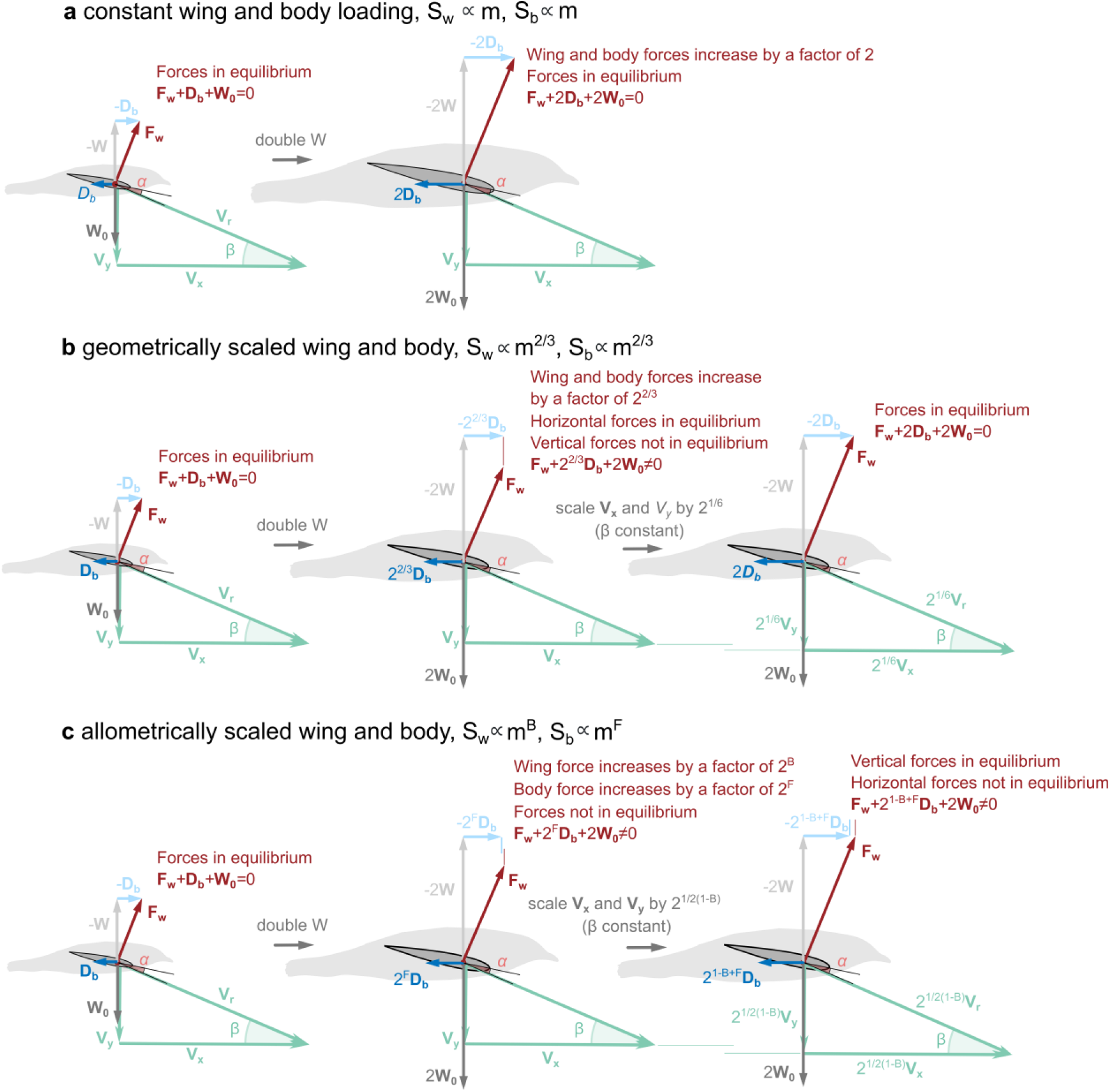
A comparison of different scaling models of wings and bodies in predictive simulations of minimum COT cruising flight. **a-c** Cartoons illustrating an example of how doubling mass influences required kinematics for cruise depending on the scaling method: constant wing and body loading systems (**a**) require no changes to kinematics; geometrically scaled wings and bodies (**b**) require flight speed and flapping speed to scale by 2^1/6^; for allometrically scaled wings and bodies (**c**) scaling the flight speed and flapping speed by 2^1/2(1-B)^, where the wing area is defined by *S*_*w*_ ∝ *m*^*B*^ and body area is defined by *S*_*w*_ ∝ *m*^*F*^, achieves weight support but does not achieve horizontal force equilibrium; flight speed and flapping speed must be scaled unequally, which leads to the change in flapping speed ratio with scale seen in Fig. 3b.

**Extended Fig. 4.**
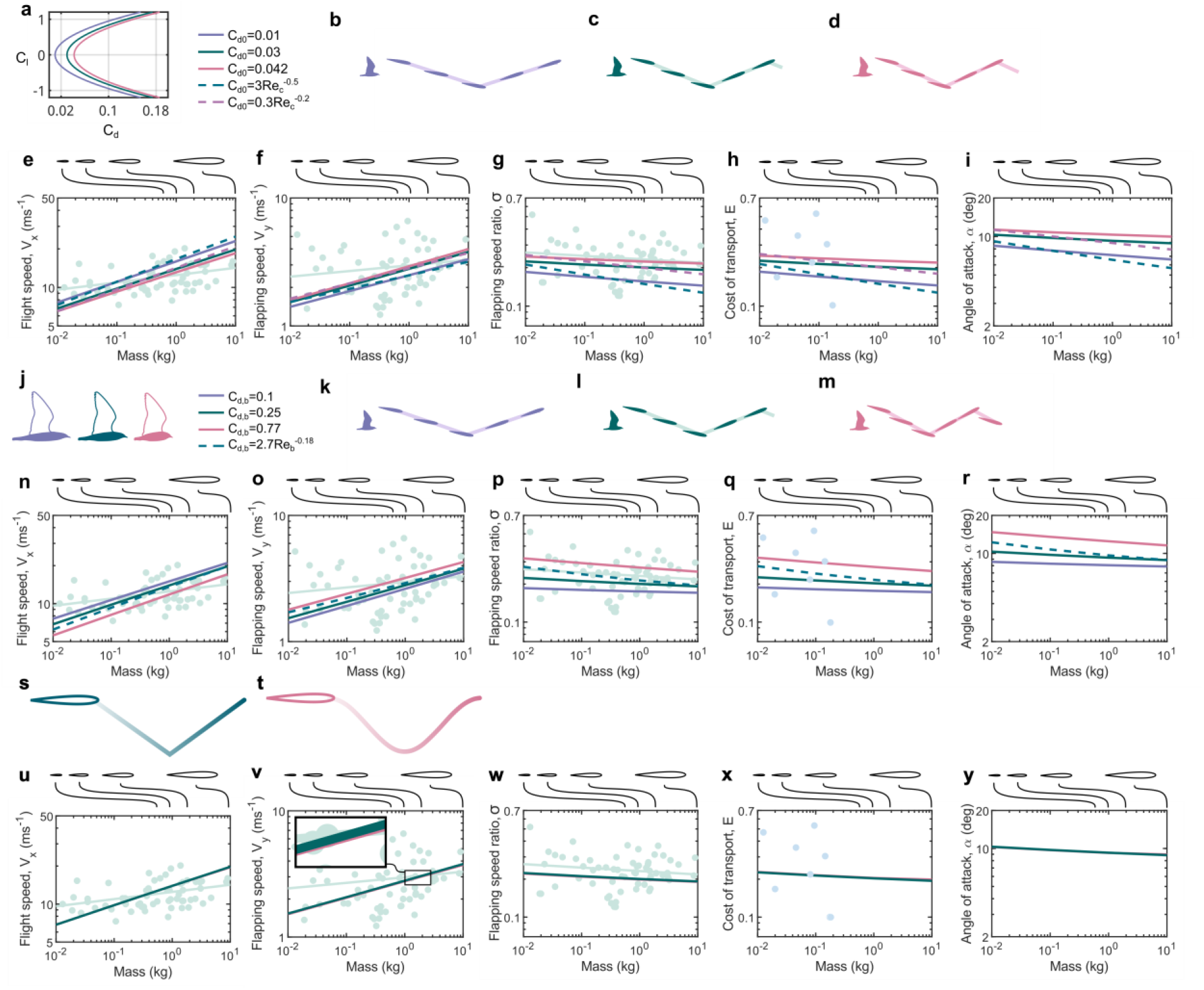
Sensitivity of the COT-optimal predicted kinematics to aerodynamic input parameters and plunge kinematic parameterisation using the two-stroke-body model. **a-i** Varying lift-independent drag coefficient. Constant drag coefficients were chosen based on the minimum value used in previous predictive simulations of bird flight^5^ (*C*_*d,0*_=0.01), and the data from wind tunnel tests on aerofoils at representative Reynolds numbers of flying birds (owl^21^: *C*_*d,0*_=0.042^5,20^). Reynolds-number dependent models of drag coefficient taken from previous predictive models 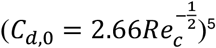 and data from wind tunnel tests on owl aerofoils (*C*_*d*,0_ = 0.33*Re*^−0.2^)^21^. **a** drag polars shown for three (constant) values of lift-independant viscous drag coefficient. **e-i** predicted COT-optimal kinematics for birds of different scales using constant (solid lines) and Reynolds-number-dependent (dashed lines) models for lift-independent drag coefficient. **j-r** Varying body parasitic drag coefficient. Constant drag coefficients were chosen based on the ranges derived from field experiments of dividing birds (0.17<*C*_*D,b*_<0.7) with an intermediate value taken from previous predictive simulations (*C*_*D,b*_ =0.25)^6^. **j** Sketches showing how body drag coefficient changes might occur due to changes in body slenderness ratio. **n-r** Predicted COT-optimal kinematics for birds of different scales using constant (solid lines) and Reynolds-number-dependent (dashed lines) models for body parasitic drag coefficient. **s-y** Comparing constant and sinusoidal flapping speeds. **s-t** sketches of wing trajectories. **u-y** Predicted COT-optimal kinematics for constant and sinusoidal flapping speed models; for the 1kg bird the mean flapping speed of the sinusoidal model is <0.01% lower than the mean flapping speed of the constant flapping speed model.

**Extended Fig. 5.**
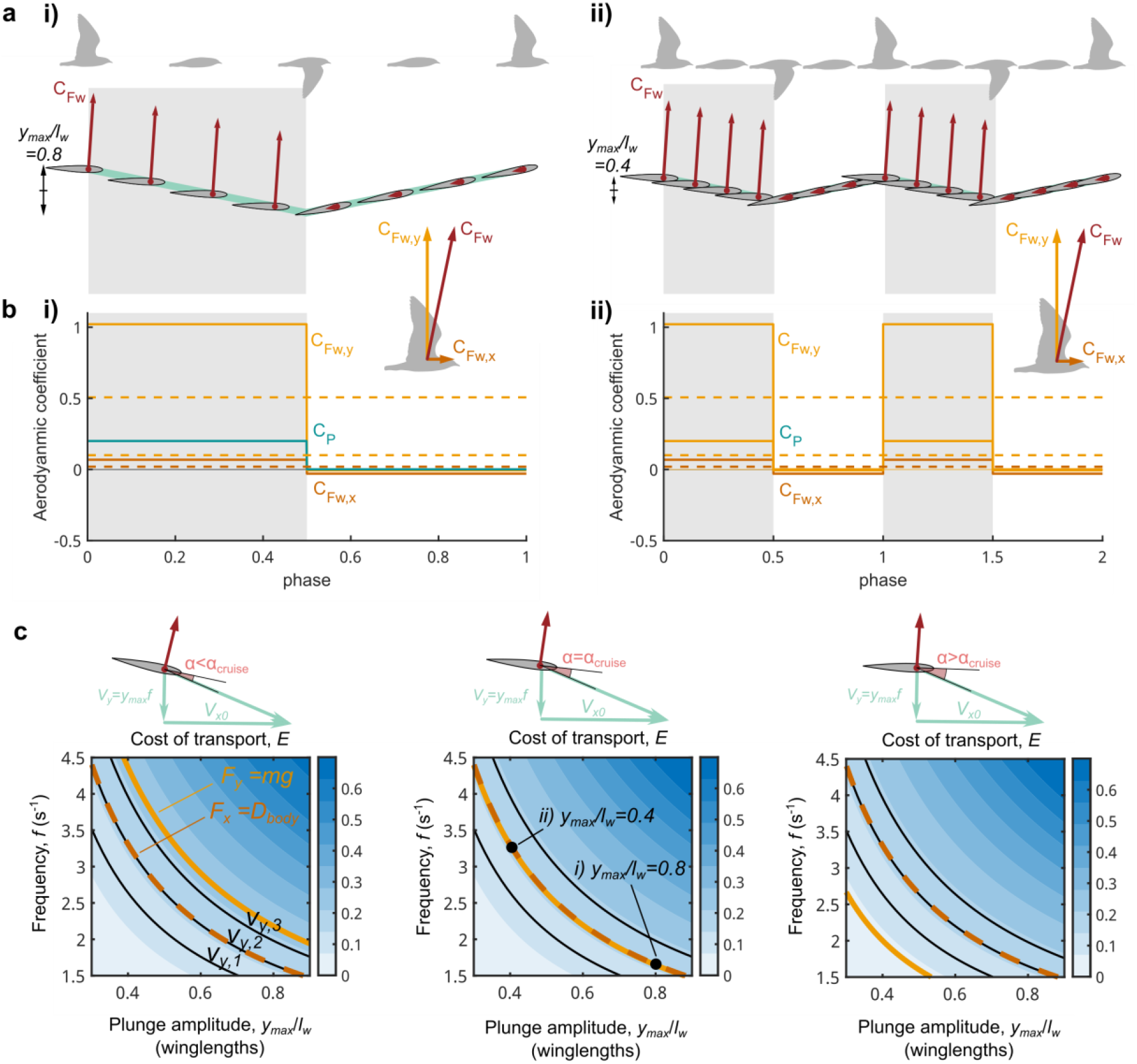
The influence of amplitude and frequency on aerodynamic force and power of plunging wings of unit span at constant flight speed, *V*_*x0*_. **a** Example kinematics of two flapping wing systems of equal, and constant flapping speeds, during the downstroke (greyed region) and upstroke (white region). Both systems have equal (positive) angles of attack on the downstroke and zero angle of attack on the upstroke. **i)** has double the amplitude and half the frequency of **ii)**. Instantaneous aerodynamic force coefficient, C_Fw_, shown by red arrows. **b** Aerodynamic force and power coefficients of the systems in **a**; solid lines are instantaneous coefficients of horizontal force (C_Fx_, dark orange) and vertical (C_Fy_, light orange) aerodynamic force, and aerodynamic power (C_P_, turquoise), and dashed lines are the respective cycle-averaged values, which are equal for **i)** and **ii). c** Contours of COT for varying plunge amplitudes, frequencies, and angles of attack. Contour of constant flapping speed have constant COT; increasing flapping speed from *V*_*y,1*_, to *V*_*y,2*_, to *V*_*y,3*_, for example, increases COT. Amplitude-frequency combinations that achieve vertical equilibrium are shown as the light orange lines, and combinations that achieve horizontal equilibrium are shown as the dashed dark orange lines. Cruise conditions are achieved where these lines overlap. At the angle of attack that achieves cruise (middle plot) there are an infinite number of amplitude-frequency combinations; points i) and ii) correspond to the combinations from **a i)** and **ii)**, respectively.

**Extended Fig. 6.**
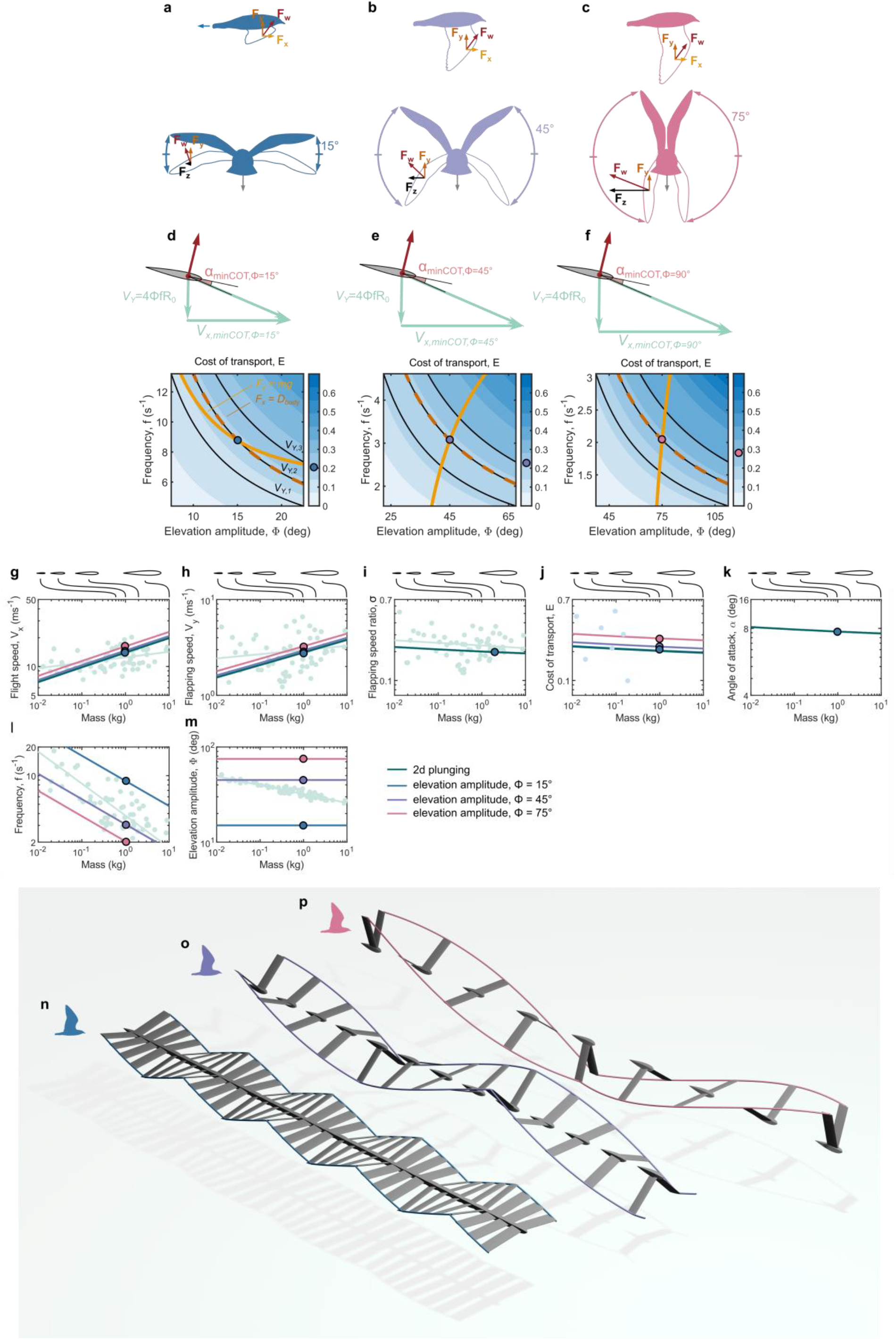
The influence of wing rotation on aerodynamics and predicted kinematics across scales of birds. **a-c** Diagrams of birds with constrained wing rotation amplitudes. Example aerodynamic force vectors are shown at the downstroke-upstroke transition. All systems have the same vertical force (*F*_*y*_) but greater amplitude systems require greater total wing aerodynamic force (*F*_*w*_) for equilibrium flight due to a component of the force being resolved laterally (*F*_*z*_); the net lateral force would be cancelled by two symmetrical flapping wings. **d-f** Contours of COT for varying elevation amplitudes and frequencies. Each plot is calculated at the flight speed and angle of attack that yielded the minimum COT solution for the particular elevation amplitude highlighted. As elevation amplitude increases the minimum COT kinematics incur higher COT. At a given flight speed and angle of attack there is only a single combination of elevation amplitude and frequency that yields force equilibrium. **g-m** Predicted trends of flight speed, COT and wing motion across scales of birds assuming fixed elevation amplitudes (Φ = 15°, blue; Φ = 45°, purple; Φ = 75°, pink), with 2d plunging model (green) shown for reference. **n-p** Visualizations of predicted kinematics for COT-optimal flight with constrained elevation amplitudes. All cases shown over equal time windows equal to the time period of a single wingbeat of the case constrained amplitude Φ = 75°.

**Extended Fig. 7.**
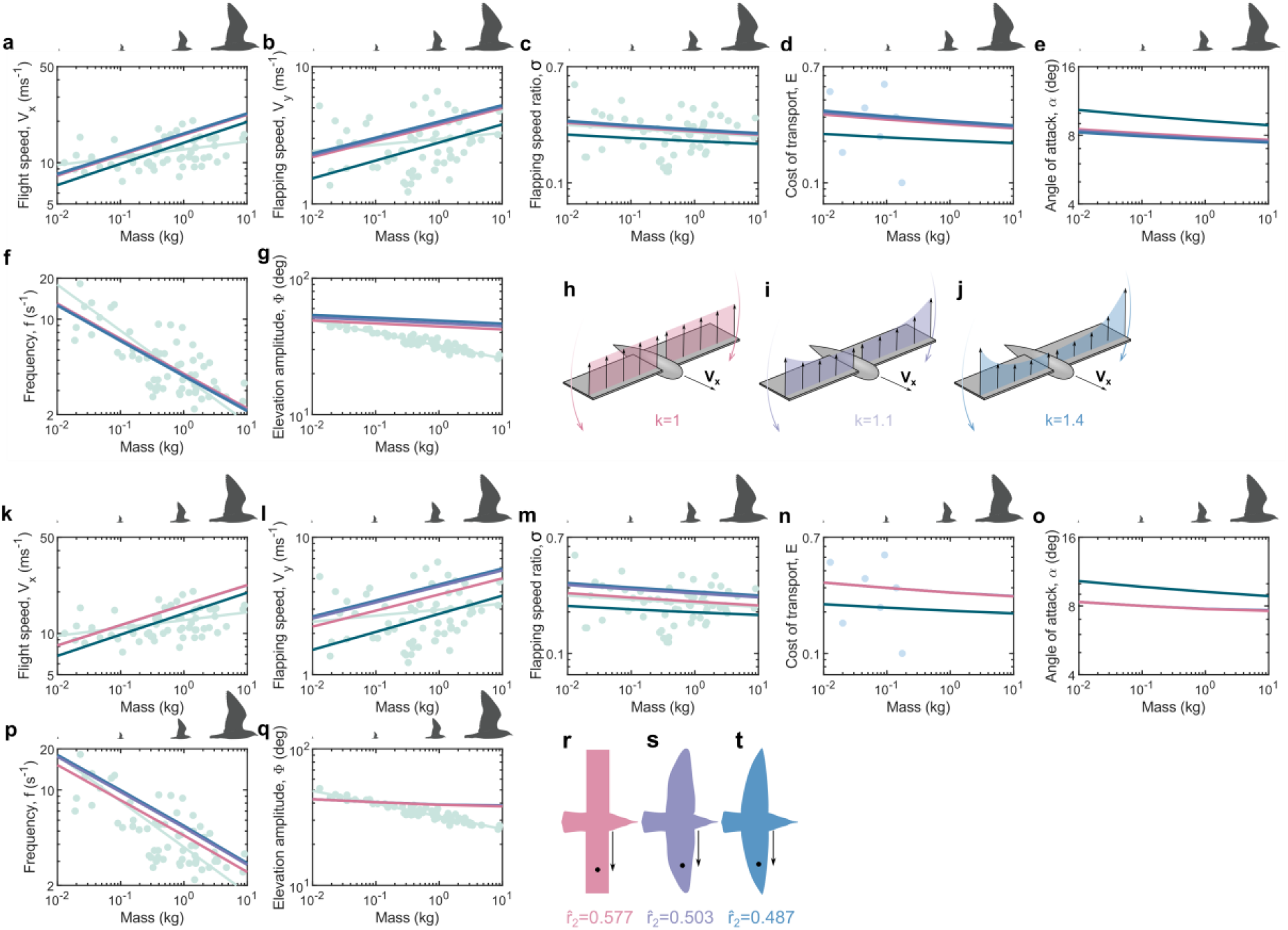
The influence of aerodynamic modelling decisions on predicted kinematics across scales of birds. **a-g** Predicted trends of COT-optimal flight speed, COT and wing motion across scales of birds with different induced power factors (*k*=1, pink; *k*=1.2, purple; *k*=1.4, blue)), with two-stroke-body model (green) shown for reference. **h-j** cartoons illustrating induced velocity distributions along the wing span for different induced power factors. **k-q** Predicted trends of COT-optimal flight speed, COT and wing motion across scales of birds with different chord distributions, characterized by the radius of the second moment of area,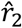), with two-stroke-body model (green) shown for reference. Examples chord distributions used were a rectangular planform (pink, 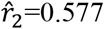), the European pied flycatcher (purple, 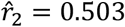)^39^, and hummingbird (blue, 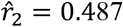)^39^, noting that for semi-elliptical planform 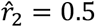. **r-t** Diagrams of wing planforms with chord distributions corresponding to the stated radii of the second moments of area.

## Supplementary Information

### Reference Frames, Kinematics and Flight Mechanics

The global reference frame (*x, y, z*) is defined with the x-axis parallel to the flight velocity vector, *V*_*x*_, the y-axis opposite to the local gravity vector, *g*, and the z-axis perpendicular to the x-y plane forming a right handed system (Extended Fig. 8). The wing elevation angle, *ϕ*, is used to described rotation from the global to the wing reference frame (*X, Y, Z*). The method of transforming between reference frames in flapping wing models is detailed elsewhere^6^.

**Extended Fig. 8.**
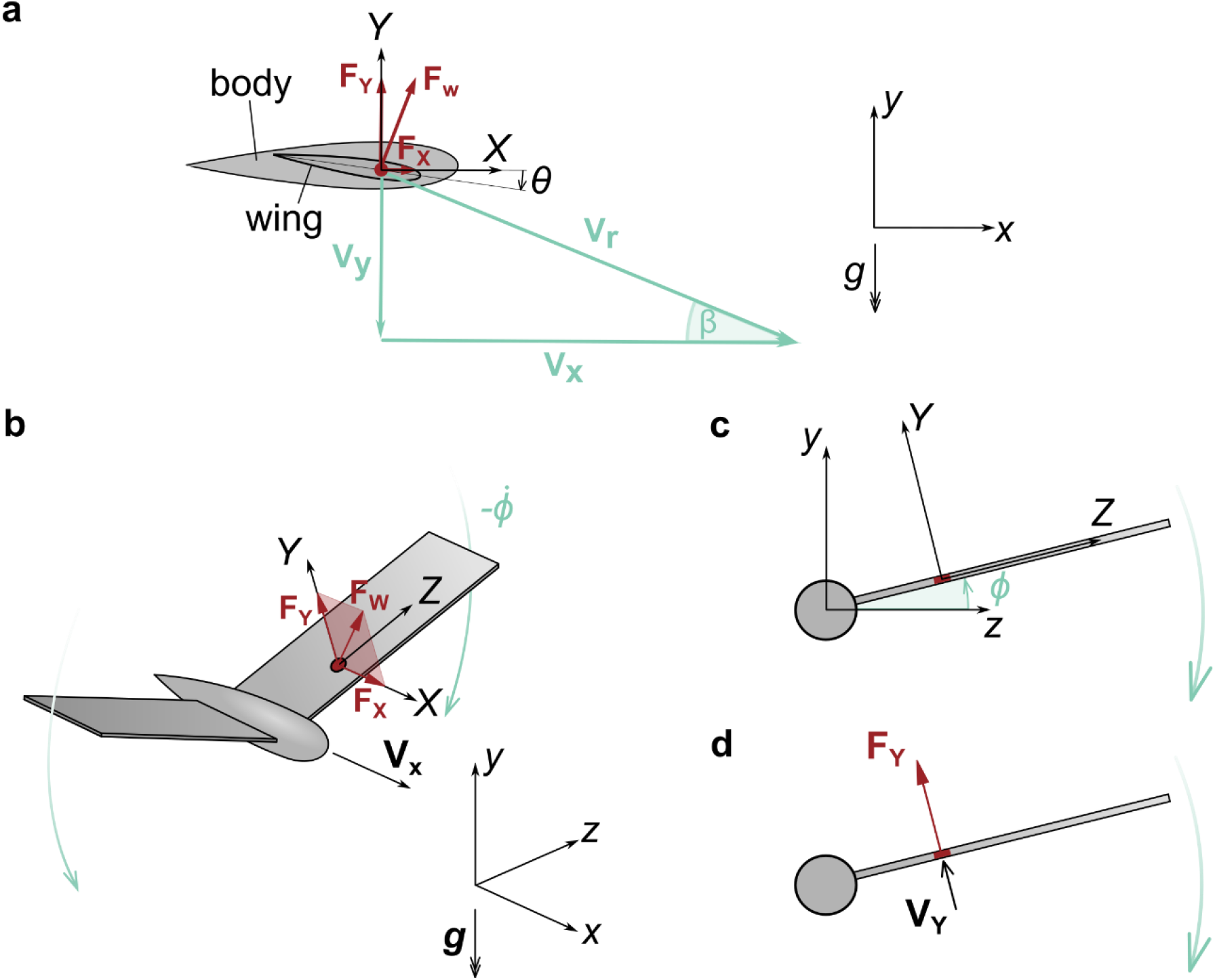
Reference frames for kinematics and force definitions. **a** Plunging wing model reference frames, with global (*x,y,z*) axes defined by the flight velocity vector, *V*_*x*_, and gravity vector, g. Wing axes (X,Y,Z) are parallel to the respective world axes, and the wing plane is the *X-Z* plane. *θ* is the wing pronation angle measured from the horizontal to the aerofoil zero-lift line. *β* is the wing plunging angle measured from the flight velocity vector to the relative wing velocity vector, V_r_. **b-d** Rotating wing model reference frames, show the wing axes rotated around the global x-axis by the wing flapping angle, *ϕ*, and the components of force and velocity normal to the wing plane, *F*_*Y*_ and *V*_*Y*_, respectively.

### Allometric Scaling Relationships

**Table 1.**

| Variable | Allometric scaling equation |
| --- | --- |
| Wing planform area, $S$ | $0.0782m^{0.718 \ 40}$ |
| Body reference area, $S_b$ | $0.0129m^{0.614 \ 41}$ |
| Wing length, $l_w$ | $0.528m^{0.396 \ 40}$ |
| Wing mass, $m_w$ | $0.0706m^{1.098 \ 40}$ |
| Wing radius of gyration, $r_0$ | $0.179m^{0.4352 \ 39}$ |

### Analytical Downstroke Model

### The Relationship Between Flight Speed and Plunging Angle

**Extended Fig. 9.**
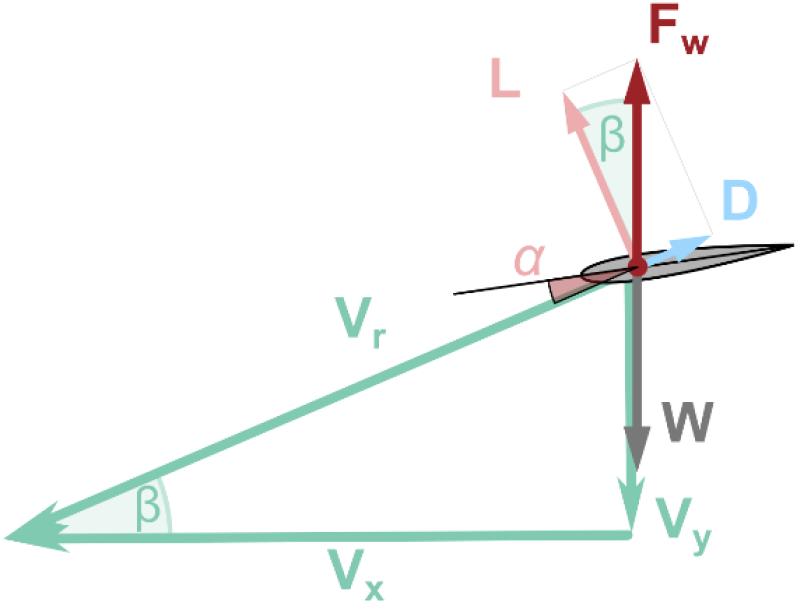
Force and velocity vector diagram for the downstroke model. The wing aerodynamic force vector, *F*_*w*_, acts only during the downstroke, and acts in the opposite direction to the weight, *W*. The aerodynamic drag, *D*, and lift, *L*, are opposite and perpendicular to the relative wing velocity, *V*_*r*_, respectively.

Consider a vertically plunging wing (Extended Fig. 9) with no induced velocity, such that 3d aerodynamic force coefficients (*C*_*D*_, *C*_*L*_) are assumed equal to the 2d aerofoil force coefficients (*C*_*d*_, *C*_*l*_). For vertical equilibrium, with a downstroke fraction of *γ*:

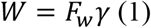

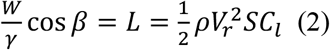

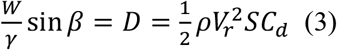

The aerofoil drag polar is given by:

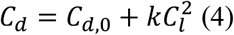

Substituting (2) and (3) into (4) gives

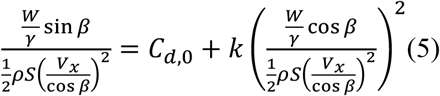

and expanding this gives

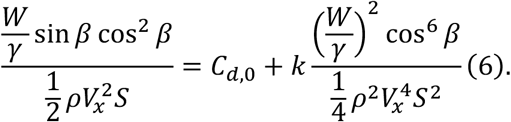

Multiply (6) by 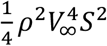 gives

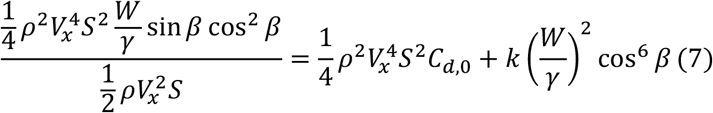

And simplifying gives

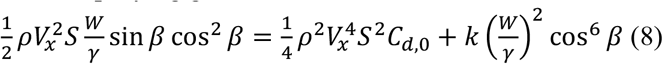

Rearrange to set terms =0:

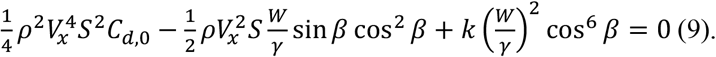

By inspection of (9) we can see that for a finite *V*_*x*_ as *β* ⟶ 0 (i.e. fixed, non-flapping wing) the left hand side of the equation is greater than 0, so the equality cannot be maintained. This is because horizontal force balance cannot be achieved by a fixed wing; whereas vertical force balance alone can be achieved by a fixed wing (equation 2).

To convert equation (9) into a quadratic let 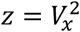:

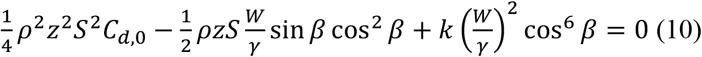

Which has the roots:

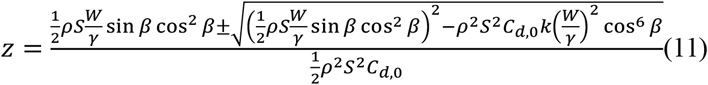

And the non-trivial solutions for flight speed:

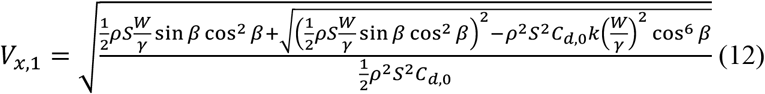

and

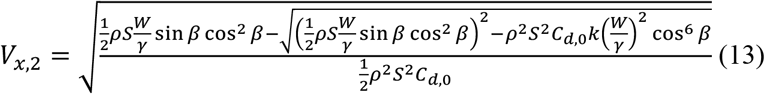

**Extended Fig. 10.**
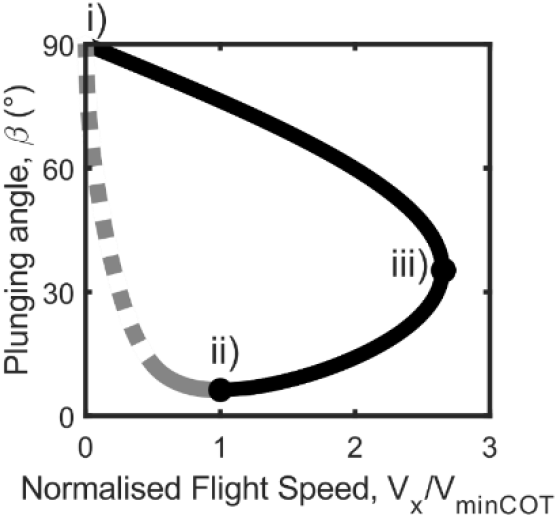
Plot of solutions of equations (12) (black line) and (13) (grey line) assuming a downstroke fraction *γ* = 0. 5. The dashed grey line show the mathematical solution to equaton 13 angles of attack where the wing would likely stall (*α* ≥ 15°) and as such are not deemed physically viable solutions for cruise. The two solution sets meet at two points on the curve. Point i) is a solution with zero flight speed and a 90° plunging angle – the wing flaps vertically downwards at zero angle of attack and achieves weight support through drag alone. Point ii), by inspection, is the solution with the smallest plunging angle. iii) represents the maximum flight speed attainable using the downstroke model.

By inspection, the minimum plunge angle (Extended Fig. 10 point ii) can be obtained by equating (12) and (13), which leads to

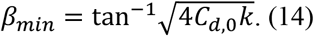

It should be noted that equation (14) is identical to the minimum glide slope of a fixed wing.

From extended Fig. 10 point ii), reducing flight speed for the *V*_*x*,2_ solutions increases the angle of attack. Below the flapping stall speed equation (4) breaks down under the assumption of a linear variation of lift coefficient and angle of attack, and equation (13) is no longer valid; for completeness the result of equation (13) is shown as a dashed line.

### Minimum Cost of Transport

The mean aerodynamic power of the downstroke is

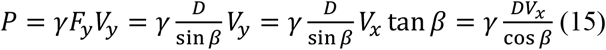

The cost of transport, E, is then given as

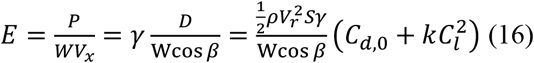

The lift coefficient is defined as

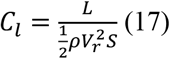

Substituting (2) into (17) gives

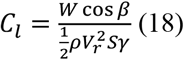

And substituting (18) into (16) gives

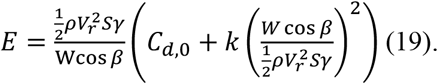

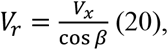

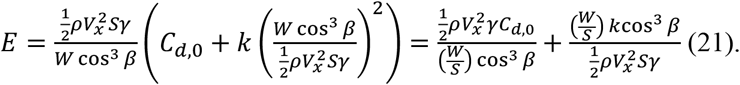

From (21) it can be seen that for very low flight speeds the second term dominates and COT can be reduced by the plunge angle tending to 90 degrees - this is the drag-based weight support mechanism. For high flight speeds the first term dominates and COT can be reduced by reducing the plunge angle, and the drag is mainly due to lift-independent viscous drag. For more typical intermediate flight speeds seen in birds the plunging angle is selected as a balance between these two cases.

Taking the partial derivate of (21) with respect to flight speed gives

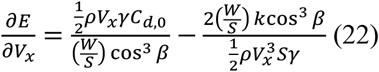

The minimum COT is defined by setting equation (22) equal to zero:

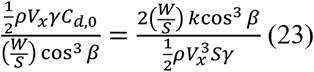

The speed at which this minimum COT occurs is then given by

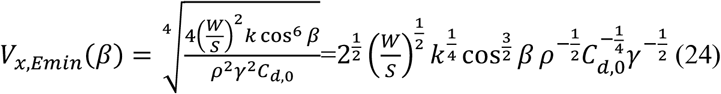

Substituting (24) into (9) gives

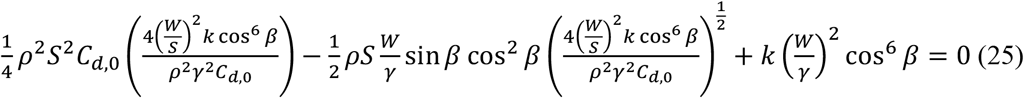

Simplifying gives

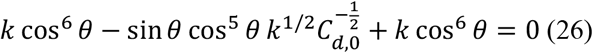

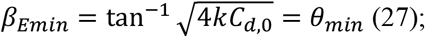

this was given as equation (1) in the main text. So the plunging angle for minimum COT is equal to the minimum plunging angle that can be used while maintaining force balance. Therefore the flight speed at which the solutions for *V*_*x*,1_ and *V*_*x*,2_ intersect is the flight speed for minimum plunging angle and for minimum COT.

Substituting equation (27) into (24) gives

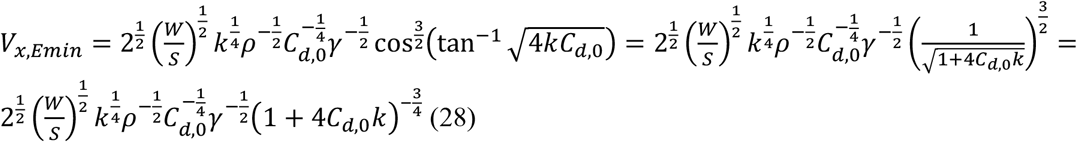

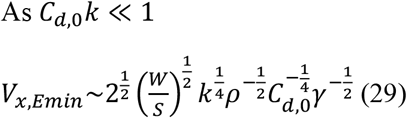

Supposing a downstroke fraction of *γ* = 1/2, The minimum COT flight speed is given as

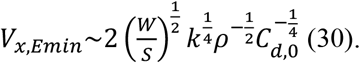

And minimum COT flapping speed is given as

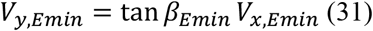

Substituting (27) and (28) into (31) gives

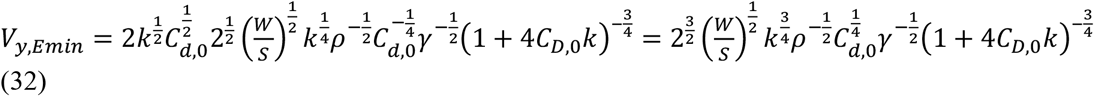

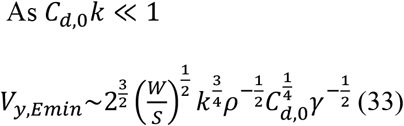

Supposing a downstroke fraction of *γ* =1/2, the minimum COT flapping speed is then

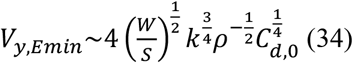

Recalling that the angle of attack is given by

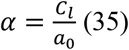

Rewriting (2) gives

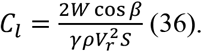

Substituting (36) and (20) into (35) gives

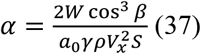

And substituting in the minimum COT plunge angle (27) and minimum COT flight speed (28) gives

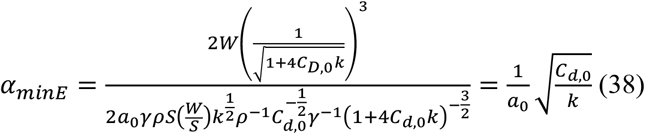

Which is identical to the angle of attack for maximum lift/drag ratio of a fixed 2d aerofoil. Note that equation (38) is independent of downstroke fraction.

Substituting the minimum COT plunge angle (27) and minimum COT flight speed (28) into (21) gives the minimum COT as

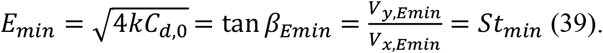

### Maximum Cruise Speed

The upper limit on flight speed that can be predicted using the downstroke model is given by considering equation (9): by inspection, as *V*_*x*_ increases the first term increases in magnitude more quickly than the second term. Therefore, there becomes a point where the positive terms are greater than the negative term and the equality can no longer be achieved.

As an approximate solution, as *V*_*x*_ becomes large the first two terms become larger than the third, so equation (9) can be simplified as

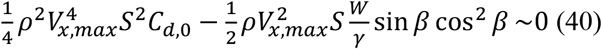

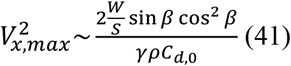

Differentiating (41) with respect to the plunging angle gives

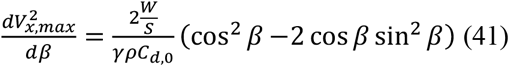

As we are only interested in positive flight speeds, the maximum flight speed can be obtained by finding the plunging angle at which the square of the flight speed is maximum:

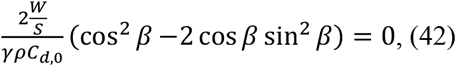

and

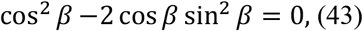

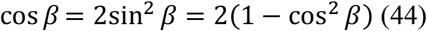

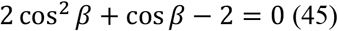

Which gives the non-trivial solution of

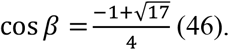

Substituting (46) into (41) and taking the square root gives the approximate solution for the maximum flight speed of the downwstroke model:

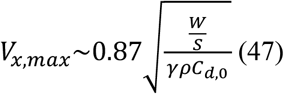

### Summary of key equations

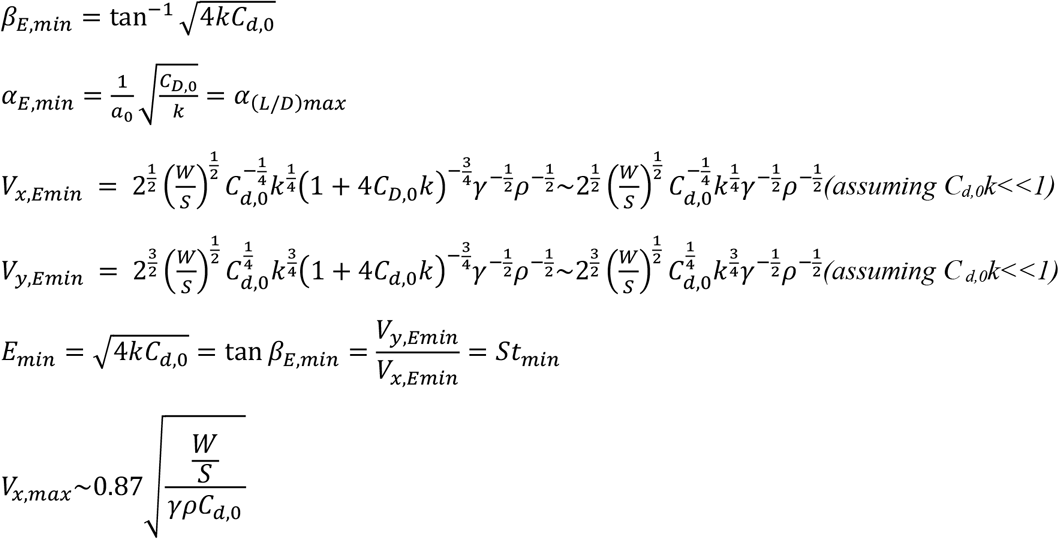

### Analytical Two-Stroke Model

Extended Fig. 11 illustrates the process of creating effective drag polars for the two-stroke and two-stroke body models, assuming a downstroke fraction, *γ* = 1/2. Minimum COT cruise conditions were found to use low plunging angles (Fig. 2). So the influence of additional drag on the upstroke or from the body can be approximated by incrementing the effective zero lift drag coefficient on the downstroke to incorporate the drag produced during the upstroke; for steeper plunging angles the accuracy of this approximation would reduce. The effective drag coefficient can be modelled by summing the drag coefficients on the downstroke and upstroke:

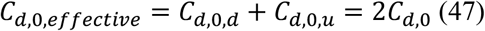

The effective zero-lift drag coefficient in equation (47) can be substituted into equations (27), (38), (29), (32) and (39) to give the minimum COT plunging angle, angle of attack, flight speed, flapping speed and COT, respectively, for the two-stroke model:

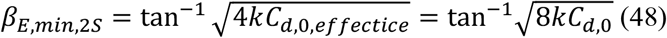

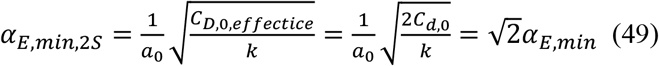

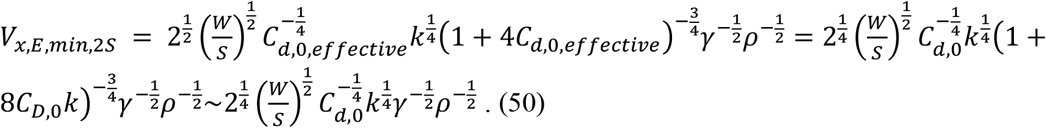

As *γ* = 1/2, the flight speed is given as

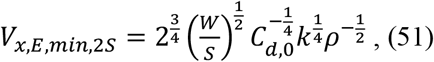

which can also be written in terms of the flight speed of the downstroke model, which is the system with the lowest possible COT

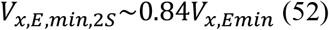

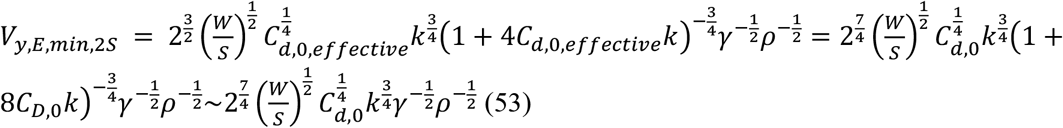

As *γ* = 1/2, the flapping speed is given as

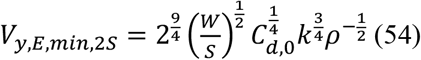

which can also be written in terms of the flapping speed of the downstroke model, which is the system with the lowest possible COT

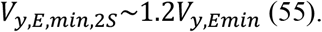

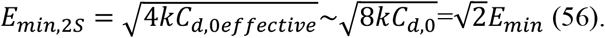

**Extended Fig. 11.**
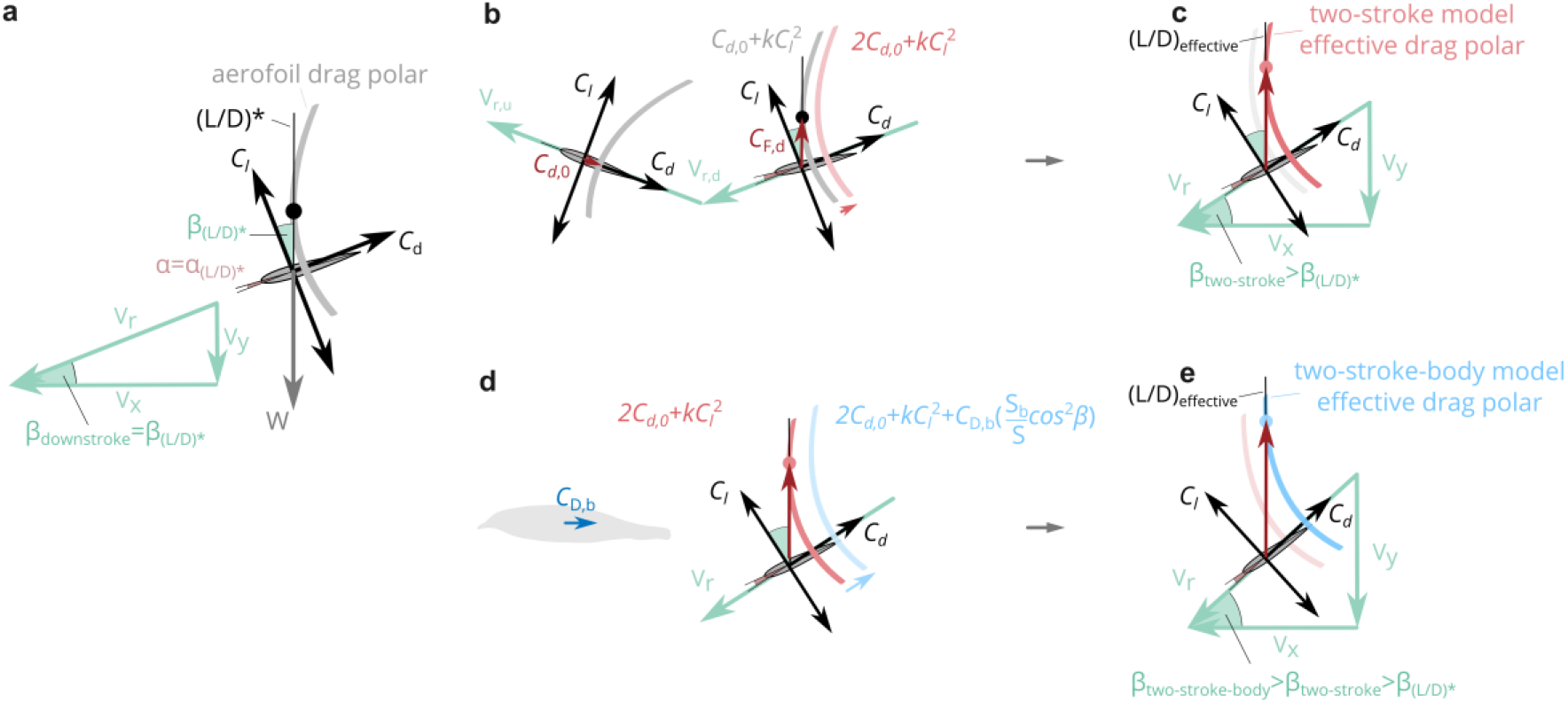
The influence of upstroke and body drag on the effective drag polar. **a** The downstroke model in minimum COT flight conditions, with flight speed, *V*_*x*_, and plunge speed, *V*_*y*_. The aerofoil drag polar is rotated by the plunging angle, 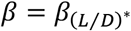, which yields the maximum lift-drag ratio, (*L*/*D*)^*\**^. The weight vector, *W*, is parallel to the tangent line to the drag polar. **b** The two-stroke model, showing how the drag on the upstroke can be incorporated into the effective zero lift drag coefficient of the downstroke. **c** The two-stroke model in minimum COT flight conditions; the effective drag polar is rotated by the plunging angle *β*_*two*−*stroke*_, which gives the maximum effective lift to drag ratio for this system. **d** The two-stroke-body model, showing how the drag on the body can be incorporated in to the effective zero lift drag coefficient the downstroke. **e** The two-stroke-body model in minimum COT flight conditions; the effective drag polar is rotated by the plunging angle, *β*_*two*−*stroke*−*body*_, which the maximum effective lift to drag ratio for this system.

### Analytical Two-Stroke-Body Model

A similar approach can be taken to include the effects of both the upstroke and the body drag. The body drag should be nondimensionalized by the same reference area and reference velocity as the wing, *S*_*w*_ and *V*_∞_, respectively. The effective zero-lift drag coefficient on the downstroke is given as

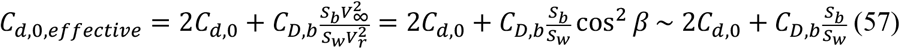

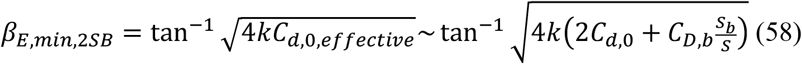

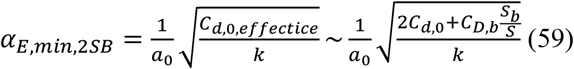

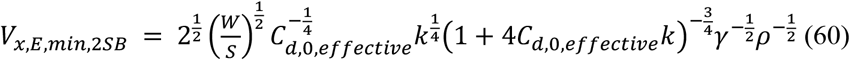

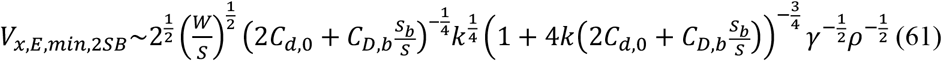

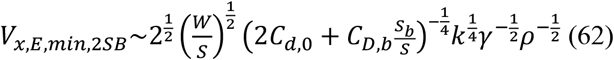

As *γ* = 1/2, the flight speed is given as

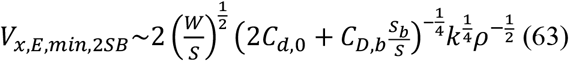

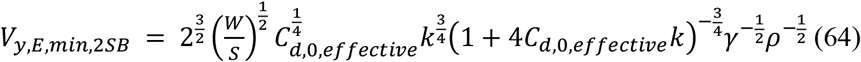

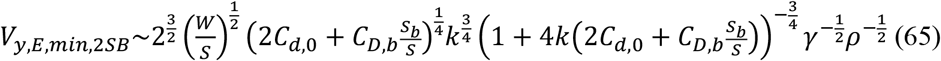

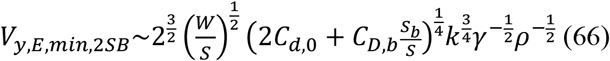

As *γ* = 1/2, the flapping speed is given as

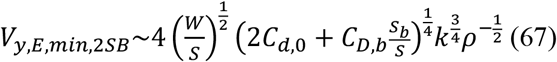

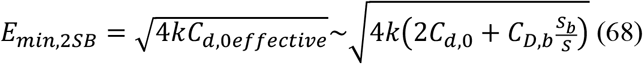

### Wing Planform Effects on Aerodynamics

The wing planform shape is described through a wing chord distribution, *c*(*r*), where *r* is the spanwise displacement measured from the shoulder joint towards the wing tip. For given kinematics the planform shape influences the induced and power drag of the wing. The induced drag is considered through the actuator disk model of induced velocity described in Methods, with the induced power factor, *k*, accounting for planform shape. This is equivalent to assuming an induced velocity correction factor^33^. The profile power is determined by considering the aerodynamic force distribution on the wing, with lift per unit span varies along the wing length as

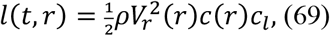

lift on an infinitesimal strip is given as

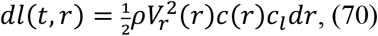

and the total lift on the wing is given as

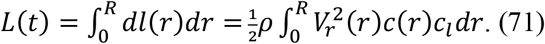

A combination of twist and change in aerofoil geometry can be used to allow each aerofoil to operate at the same lift coefficient (e.g. one that gives optimal aerodynamic efficient). This assumption is made here, leading to

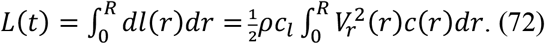

For a wing rotating around a single axis at the shoulder joint the relative wind velocity is given by

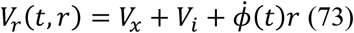

where 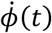 is the angular velocity. This leads to the lift being given as

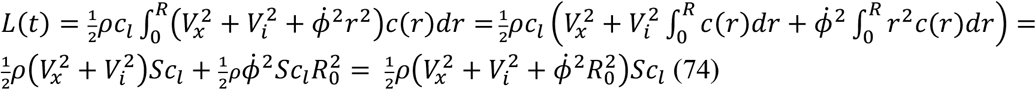

where *S* is the wing planform area, and *R*_0_ is the radius of the second moment of area of the wing, commonly written in nondimensional form as

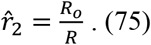

The term 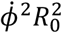 in (53) is the square of the tangential velocity due to wing rotation of an aerodynamic control point located at a distance R0 in the spanwise direction from the shoulder joint; 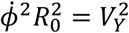.

An equivalent equation to (53) can be written for the drag:

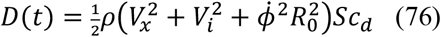

These expressions are used to account for the influence of wing planform geometry on profile lift, drag and power, by adjusting the value of the nondimensional radius of second moment of area 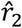. This value has been described elsewhere for some birds 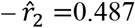 (European pied flycatcher^39^), 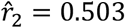 (hummingbird^39^); for a wing with a uniform chord distribution 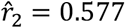. The sensitivity of the predicted kinematics and COT to 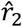 is shown in Extended Fig. 7.

